# Topographic and somatotopic organization of functional connectivity between the intraparietal sulcus and primary somatosensory cortex in macaque monkeys

**DOI:** 10.64898/2026.09.19.752858

**Authors:** Wei-An Sheng, Simon Clavagnier, Mathilda Froesel, Tobias Heed, Wim Vanduffel, Suliann Ben Hamed

## Abstract

The posterior parietal cortex integrates somatosensory information with signals supporting action, spatial cognition, and multisensory processing, yet the fine-scale topographic relationship between the intraparietal sulcus (IPS) and primary somatosensory cortex (S1) remains poorly understood. Here, we used resting-state fMRI in 10 awake rhesus macaques to characterize functional connectivity between four S1 subregions (areas 3a, 3b, 1, and 2) and 223 single-voxel seeds densely sampling an IPS-centered parietal territory. Across S1 subregions and hemispheres, IPS-S1 connectivity exhibited a highly structured spatial organization characterized by systematic anteroposterior and mediolateral gradients and a prominent semi-oval topology following the geometry of the intraparietal sulcus. Connectivity was generally stronger toward the cortical banks and convexities and weaker near the sulcal fundus, a pattern that could not be explained by reduced local temporal signal-to-noise ratio. Unsupervised hierarchical clustering independently recovered this spatial organization and identified connectivity-defined subdivisions that only partially corresponded to classical cytoarchitectonic boundaries. Along the S1 mediolateral axis, prominent troughs in cluster-averaged connectivity profiles corresponded to several previously described somatotopic transitions, allowing to consistently map putative boundaries between tongue and face representations and between hand and trunk-leg representations. Based on these boundaries, individual IPS clusters exhibited connectivity with multiple S1 body-part sectors, with the pattern of connectivity across these sectors varying across IPS clusters and S1 subregions. Together, our findings reveal that IPS-S1 functional connectivity is organized by a large-scale sulcal topology while preserving fine-grained features of S1 somatotopic organization. These findings suggest that the functional organization of the IPS is shaped by multiple spatial dimensions, including large-scale sulcal topology and the organization of S1 subregions and body-part representations, which may together contribute to the functional specialization of individual IPS regions.

## Introduction

The posterior parietal cortex (PPC) occupies a central position within the cortical sensorimotor network, serving as an interface between primary sensory areas with higher-order systems involved in spatial cognition and action planning (Andersen et al., 1997; Freedman and Ibos, 2018; Medendorp and Heed, 2019). Numerous PPC regions receive somatosensory input and combine it with visual, proprioceptive, and motor-related signals to support functions such as body-centered representations, spatial perception, reaching, grasping, and the control of movements in peripersonal space (Andersen and Buneo, 2002; Graziano and Cooke, 2006; Andersen and Cui, 2009). These computations depend critically on interactions between PPC and the primary somatosensory cortex (S1), which provides detailed information about tactile stimulation and body posture (Jones and Powell, 1970; Delhaye et al., 2018).

Within the PPC, the intraparietal sulcus (IPS) is a key hub for integrating multisensory information that receives substantial somatosensory input. The IPS contains many of the principal sensorimotor and multisensory PPC areas, which can be broadly grouped into regions supporting somatosensory and motor integration (areas 5, 7b), visuomotor transformations for reaching and grasping (MIP, AIP), spatial attention and oculomotor control (LIP, 7a), and multisensory encoding of peripersonal space (VIP). In particular, the ventral intraparietal area (VIP) exhibits sensitivity to tactile and proprioceptive signals, supporting functions ranging from body representation to sensorimotor transformations (Duhamel et al., 1998; Avillac et al., 2005; Niu et al., 2020; for review, Foster*, Sheng*, et al. 2022). Neurons within the fundus of the IPS show robust responses to tactile, visual, and auditory stimulation, often with receptive fields centered on the face, highlighting a specialization for encoding peripersonal space and integrating somatosensory signals with higher-order spatial and action-related processes (Cooke and Graziano, 2003; Guipponi et al., 2013; Sheng et al., 2025). Crucially, these areas are not organized as discrete functional islands but maintain a continuous topographic layout along both the anteroposterior and mediolateral axes of the sulcus (Hyvärinen, 1981; Andersen et al., 1990; Snyder et al., 1997; Duhamel et al., 1998; Murata et al., 2000). Such gradient-like, large-scale organization raises the question of whether IPS-S1 functional connectivity likewise varies systematically across IPS space, rather than changing only at discrete areal boundaries.

The primary somatosensory cortex (S1), in contrast, is classically described according to its somatotopic organization, with representations of different body parts systematically arranged along the mediolateral axis of the postcentral gyrus (Penfield and Boldrey, 1937; Kaas, 1983). Although this organization is often described in terms of distinct body-part territories, neighboring representations are embedded within a continuous cortical map of the body surface. This raises an important question for understanding IPS-S1 interactions: how are transitions between neighboring somatotopic representations reflected in large-scale functional connectivity? This orderly organization provides a structured input space through which tactile and proprioceptive information is encoded and relayed to higher-order cortical areas. Despite extensive knowledge of S1 organization and IPS function, the precise topographical relationship between S1 and IPS, specifically, how somatotopically organized signals in S1 are functionally linked to distinct IPS subregions in the primate brain remains poorly understood. In particular, it is unknown whether the continuous spatial organization observed across the IPS is related to systematic changes across the S1 somatotopic map and whether transitions between classical body-part representations are reflected in IPS-S1 functional connectivity.

Anatomical tracing, electrophysiological recordings, and functional imaging studies have demonstrated extensive interactions between PPC and S1 and have characterized the connectivity, functional properties, and somatotopic organization of individual PPC and S1 cortical areas (Jones et al., 1978; Lewis and Van Essen, 2000a; Rozzi et al., 2006; Arcaro et al., 2019). However, these approaches have typically focused on predefined regions of interest, leaving unresolved how functional connectivity varies continuously across PPC space and whether transitions in connectivity correspond to classical areal boundaries or to the somatotopic organization of S1. Indeed, conventional area-based approaches collapse connectivity within predefined areal boundaries, may obscure gradients and transitions that occur within or between areas. Systematic single-voxel sampling across the full IPS extent instead allows functional connectivity to be examined as a spatially continuous phenomenon, allowing its organization to be read out directly from the data without imposing a priori areal constraints (Haak et al., 2018; Sheng et al., 2025). Functional connectivity further complements traditional anatomical and physiological approaches by characterizing relationships among distributed cortical regions and has proven useful for identifying connectivity-defined functional subdivisions in macaque cortex (Biswal et al., 1995; Fox and Raichle, 2007; Hutchison and Everling, 2014).

Previous studies suggest that IPS subregions exhibit structured connectivity with somatosensory cortex and may preferentially interact with specific body-part representations in S1 (Jones et al., 1978; Taoka et al., 1998; Gharbawie et al., 2011; Huang et al., 2012; Seelke et al., 2012), but a systematic characterization of this connectivity at a fine spatial scale is lacking. In the present study, we used resting-state functional connectivity to characterize interactions between S1 and an IPS-centered PPC territory in macaque monkeys. We asked three related questions: (1) how IPS-S1 connectivity varies as a function of position within both PPC and S1; (2) whether connectivity-defined transitions correspond to established cytoarchitectonic boundaries within PPC; and (3) whether systematic changes in IPS-S1 connectivity correspond to transitions between neighboring somatotopic representations within S1. We hypothesized that distinct portions of the IPS-centered PPC territory would exhibit different connectivity relationships with S1, giving rise to systematic spatial gradients, connectivity-defined subdivisions, and transitions corresponding to somatotopic organization.

## Material and Methods

We reused data from previous studies of Wim Vanduffel’s lab (Mantini et al., 2011, 2013; Pijnenburg et al., 2019). These data were produced in different experiments; however, the recording parameters relevant for the purposes and analyses of the present paper were similar.

### 1. Subjects

10 rhesus monkeys (Macaca mulatta) participated in the study (6 females, 4 males). They sat in a sphinx position inside a plastic primate chair (Vanduffel et al., 2001), with their heads constrained by a plastic headpost. The monkeys were trained to maintain fixation at a small red fixation point on a uniform gray background within a 2° × 3° virtual window in the center of the screen, while eye movements were monitored by a pupil-corneal reflection eye tracking system (ISCAN, RK-726PCI) at 60 Hz. The monkeys were rewarded for correct eye fixation behavior, reward frequency increasing to a threshold for longer fixations. Before each scanning session, 8-11 mg/kg monocrystalline iron oxide nanoparticle (Molday ION, BioPAL) was injected via the femoral/saphenous vein to improve the contrast-to-noise ratio (CNR) and to avoid the contribution of superficial draining veins (Vanduffel et al., 2001; Leite et al., 2002). Animal care and experimental procedures were performed in accordance with the National Institute of Health’s Guide for the Care and Use of Laboratory Animals, the European legislation (Directive 2010/63/EU) and were approved by the Animal Ethics Committee of the KU Leuven.

### 2. (f)MRI acquisition

The in-vivo MRI scans were performed with a 3T MR Siemens Trio scanner in Leuven, Belgium. For functional measurements, gradient echo planar images (GE EPI) covering the whole brain were acquired with an eight-channel phased-array receive coil and a saddle-shaped, radial transmit-only surface coil (see Kolster et al., 2014; 40 slices; 84-by-84 in-plane; flip angle=75°; repetition time (TR) = 2.0 s or 1.4s depending on the monkey; echo time (TE) = 19 ms; voxel size = 1.25 mm x 1.25 mm x 1.25 mm voxels). T1-weighted anatomical images were obtained during different sessions using a magnetization-prepared rapid gradient echo (MP-RAGE) sequence (TR = 2200 ms, TE = 4.06 ms, voxel size = 0.5 mm x 0.5 mm x 0.5 mm). During the anatomical scans, the animals were sedated using ketamine/xylazine (ketamine 10 mg/kg I.M., xylazine 0.5 mg/kg I.M., maintenance dose of 0.01-0.05 mg ketamine per minute I.V.). The specific acquisition parameters varied across studies and animals (see Table 1 in Sheng et al., 2025, longer runs and higher number of runs were acquired for monkeys achieving better fixation). This resulted in a more robust evaluation of functional connectivity in these individuals. The shortest cumulated resting-state scanning duration (81 min), the shortest duration of individual runs (10 min), and the lowest number of runs (8 runs) combined is in the standard recommendations in the field (Birn et al., 2013). The variability induced by the differences in scanning parameters is thus expected to be minimal, and if anything, it is expected to enhance the robustness of our results, as our approach highlights common connectivity patterns across all monkeys, irrespective of low-level methodological differences.

### 3. Data analysis

#### 3.1 Preprocessing

Functional time-series were realigned, coregistered to the anatomical images and normalized to F99 macaque brain template with 1mm isotropic resolution (Van Essen, 2004). The time-series were band-pass filtered (0.01–0.1 Hz or 0.0025–0.05 Hz depending on monkey), nuisance corrected for ventricle and white matter signal and motion-scrubbed for movement artefacts within and across sessions and slice time, both by means of regression analysis (Vincent et al., 2007). A spatial smoothing was then applied with a 1.5mm Full Width at Half Maximum (FWHM) Gaussian Kernel.

#### 3.2 Region of interest (ROI)

The aim of this study was to delineate the detailed inter-areal functional connectivity between the intraparietal sulcus (IPS) and primary somatosensory cortex (S1) using single-voxel seeds. To achieve this, we first manually selected 16 consecutive coronal slices from the F99 T1 anatomical volume along the anteroposterior axis to cover the full extent of the IPS (Fig. 1A). Within each coronal slice, five to seven single-voxel seeds were defined across the medial and lateral banks of the IPS using a non-uniform sampling strategy adapted to the local geometry of the sulcus (Fig. 1B, D). Seeds near the fundus were placed in adjacent voxels to provide higher spatial sampling within this narrow region, whereas seeds closer to the cortical convexity were sampled from one of three neighboring voxels because the cortical territories along the banks were broader. Consequently, inter-seed distances were shorter near the fundus than near the cortical convexities.This procedure yielded a total of 223 seeds distributed across several parietal areas, including areas 2, 2Ve, 5, 7a, 7b, MIP, VIP, PIP, V3A, AIP, and LIP (Fig. 1B), according to the Markov atlas (Markov et al., 2014).

**Figure 1.**
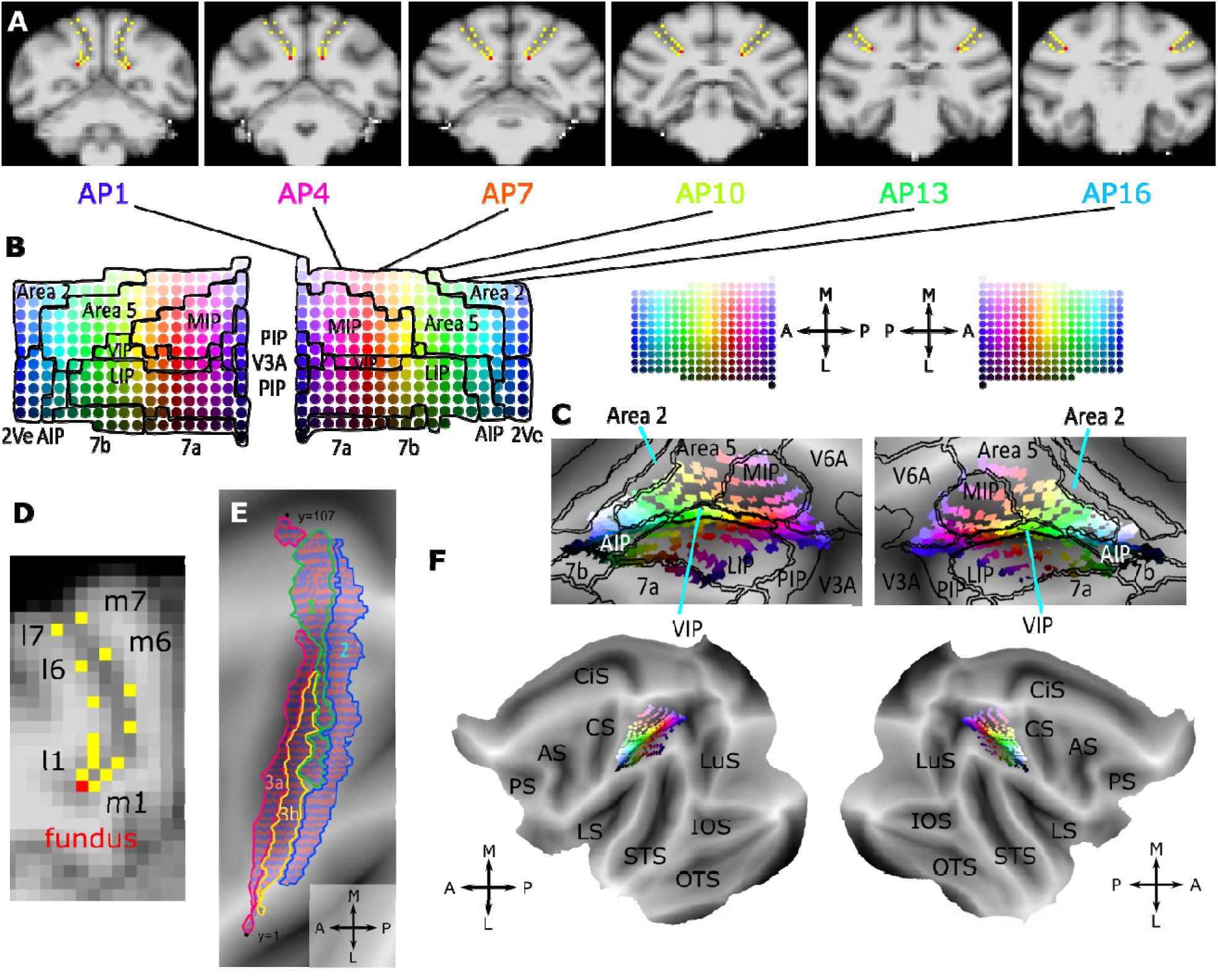
Intraparietal sulcus (IPS) seeds and sampling of their functional connectivity in the primary somatosensory area (S1). **A.** Examples of six coronal slices in the volumetric space showing the seeds along the IPS, with the anteroposterior positions indicated in B. **B.** Distribution of the 223 seeds across cortical areas, as defined by the Markov atlas (Markov et al., 2014), shown for both hemispheres. Seed locations are color-coded such that hue represents position along the anterior-posterior axis, while variations in value (lightness) and chroma (saturation) distinguish positions across the medial and lateral banks of the IPS. AIP, anterior intraparietal area; LIP, lateral intraparietal area; MIP, medial intraparietal area; PIP, posterior intraparietal area; VIP, ventral intraparietal area. **C.** Projection of seeds on flatmaps of both hemispheres, colored as in B. Note that projection of the seeds on flatmap is only for display. Functional connectivity is calculated in volumetric space. **D.** Magnified view of AP1 in A, showing seed distribution across the medial and lateral banks of the left IPS. The fundus seed is highlighted in red, whereas the remaining seeds are shown in yellow. Seeds near the fundus are placed in adjacent voxels, while those closer to the cortical convexity are sampled from one of three neighboring voxels. **E.** Identification of areas 3a (magenta), 3b (yellow), 1 (green), and 2 (blue) in the left hemisphere from which functional connectivity to IPS is computed. Note that the mediolateral extent of areas 3a and 3b is shorter than that of areas 1 and 2, reflecting the burial of these subregions within the posterior bank of the central sulcus and the consequently reduced number of surface voxels accessible at the volumetric resolution of the fMRI data. To reduce data dimensionality, surface voxels sharing the same mediolateral coordinate were averaged (see Methods 3.3). Red-blue stripes indicate successive mediolateral levels of surface voxels across all S1 subregions. A, anterior; P, posterior; M, medial; L, lateral, represent brain orientations. **F.** Same as C on the whole brain of both hemispheres. AS, arcuate sulcus; CiS, cingulate sulcus; CS, central sulcus; IOS, inferior occipital sulcus; LS, lateral sulcus; LuS, lunate sulcus; OTS, occipitotemporal sulcus; PS, precentral sulcus; STS, superior temporal sulcus

### 3.3 Functional connectivity

#### 3.3.1 Seed-based functional connectivity and S1 dimensionality reduction

Analyses were performed using AFNI (Cox, 1996) and FSL (Jenkinson et al., 2012 http://fsl.fmrib.ox.ac.uk/fsl/fslwiki/), implemented within MATLAB. Seed-to-whole-brain functional connectivity was computed using 223 single-voxel seeds located in the intraparietal sulcus (IPS). For each run and each monkey, connectivity was estimated by calculating the correlation between the time series of each seed and every other voxel in the brain. The resulting correlation coefficients were transformed to z-scores using Fisher’s r-to-z transformation.

For each seed, a one-sample t-test against zero was performed across runs using AFNI, separately for each monkey. This procedure yielded one z-score functional connectivity map per IPS seed per monkey. The resulting maps were thresholded at p < 0.001 (Froesel et al., 2024) to identify voxels showing significant functional connectivity.

Each individual functional connectivity map was projected from volumetric space onto the cortical surface using Connectome Workbench (Marcus et al., 2013). Owing to the properties of the projection algorithm, a single volumetric voxel could map onto one or multiple surface vertices (or faces) in the 2D flatmap (Fig. 1C, F). Similar procedures for connectivity estimation and surface-based visualization have been described previously for this dataset (Sheng et al., 2025).

We then examined functional connectivity between IPS seeds and four subregions of the primary somatosensory cortex (S1), areas 3a, 3b, 1, and 2, defined according to the Paxinos atlas (Paxinos et al., 1999). Within each S1 subregion, the number of surface voxels sampled along the anteroposterior dimension was limited, whereas the mediolateral dimension provided substantially greater spatial extent and resolution. Because the known somatotopic progression across S1 is primarily captured along this mediolateral dimension, we reduced the dimensionality of the connectivity data by averaging z-scores across surface voxels sharing the same mediolateral coordinate (Fig. 1E), thereby collapsing the anteroposterior dimension while retaining connectivity as a function of mediolateral S1 position. This procedure yielded four vectors representing the connectivity profiles as a function of S1 position for each IPS seed.

Subsequent IPS-space analyses (sliding-window and clustering) were performed on binarized representations of these thresholded connectivity maps. For each monkey and IPS seed, each S1 mediolateral position was coded as 1 when significant positive functional connectivity was detected (p < 0.001, one-sided) and 0 otherwise. We chose this analysis strategy to emphasize the spatial consistency and topology of IPS-S1 connectivity patterns while reducing sensitivity to variability in absolute connectivity magnitude across seeds and monkeys. Because the resulting dataset remained three-dimensional – given that we had reduced S1, but not IPS dimensionality – we applied two complementary dimensionality-reduction strategies.

#### 3.3.2 Spatial characterization of IPS-S1 connectivity

First, we reduced the mediolateral resolution of S1 using a sliding-window approach, decreasing the number of surface voxels from over 100 (left hemisphere: 107; right hemisphere: 103) to approximately 15 (left: 15; right: 14; Fig. 2C). Within each window, the binarized connectivity values were averaged across the included S1 positions, yielding a proportion of positions showing significant positive connectivity. This averaging strategy allowed visualization of IPS-S1 connectivity as a colormap in IPS space (Fig. 2B), illustrating how connectivity varies as a function of mediolateral position in S1 (Fig. 2A, D).

**Figure 2.**
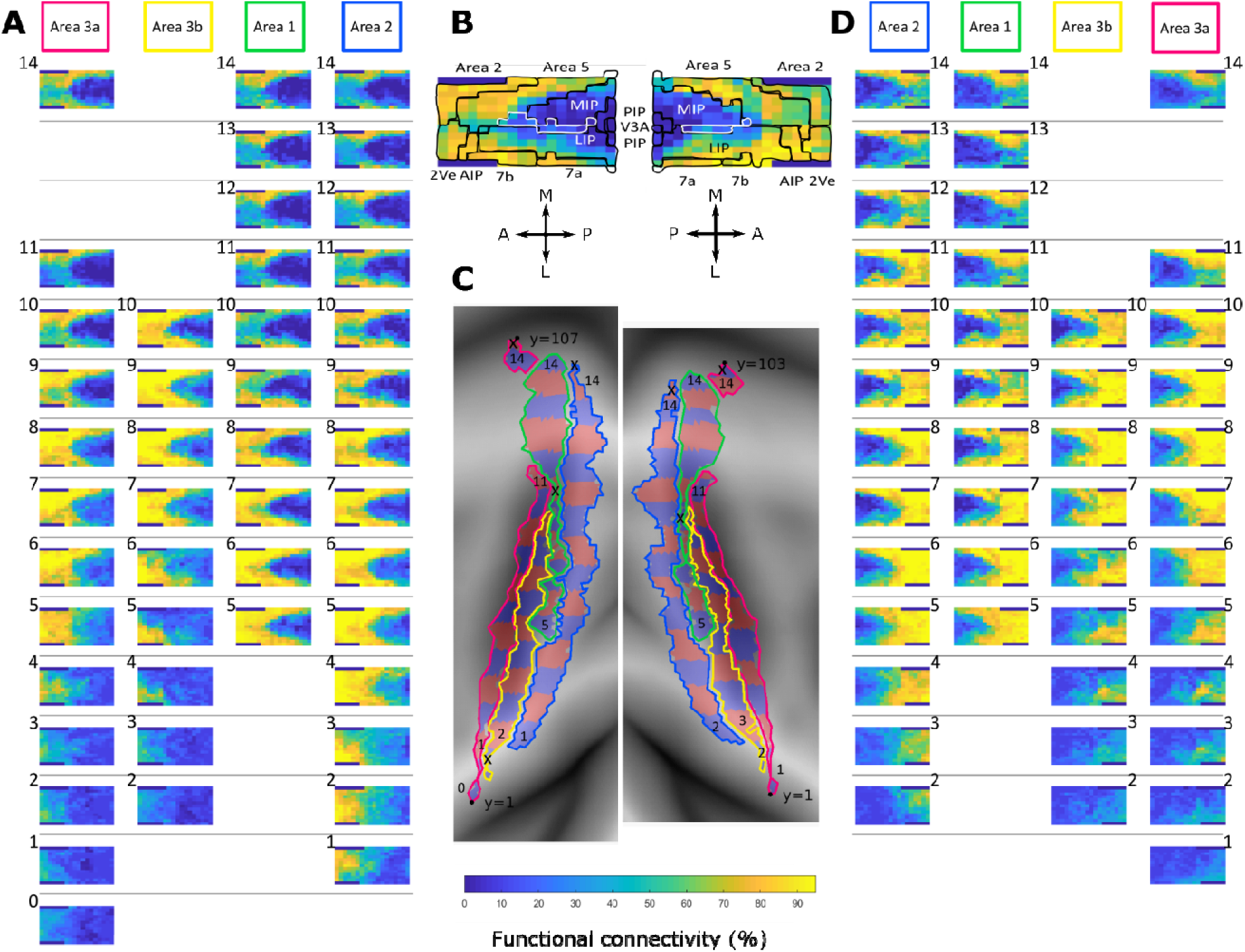
IPS functional connectivity patterns with the four S1 subregions 3a, 3b, 1 and 2 revealed by a sliding-window method. **A.** IPS-S1 functional connectivity patterns in the left hemisphere, displayed as colormaps positioned according to the spatial locations of IPS seeds. Each colormap represents the spatial distribution of positive functional connectivity between IPS seeds and one sliding-window region within a given S1 subregion, represented as binarized maps (see red-blue stripes in C). The four S1 subregions, areas 3a, 3b, 1, and 2, are arranged according to their anatomical order from area 3a, anteriorly to area 2, posteriorly. Fourteen vertical positions correspond to successive mediolateral sliding-window locations within the respective S1 subregions. The sliding-window positions are shown separately for each S1 subregion to account for differences in their surface-voxel distributions. Sliding windows containing very small surface areas are omitted. For visualization purposes, the spacing between discontinuous portions of area 3a is reduced from three units to two. Note that the F99 template is asymmetric across hemispheres; consequently, fewer mediolateral sliding windows are present in the right hemisphere than in the left. **B.** Spatial distribution of IPS seeds, with the Markov atlas areal boundaries overlaid on a representative IPS connectivity map to illustrate the anatomical location of the connectivity pattern relative to the IPS areal boundaries (Markov et al., 2014). For clarity, the two seeds located at the most medial and most lateral positions of the posterior IPS are not shown. A, anterior; P, posterior; M, medial; L, lateral. AIP, anterior intraparietal area; LIP, lateral intraparietal area; MIP, medial intraparietal area; PIP, posterior intraparietal area; VIP, ventral intraparietal area. **C.** Sliding-window regions across the four S1 subregions. Successive mediolateral window positions are indicated by alternating red-blue stripes. **D.** Same as in A, for the right hemisphere. The anatomical order of S1 subregions is mirrored relative to the left hemisphere (see schematic in B).

Second, we reduced the dimensionality of the IPS seed space by seed groups, defined as sets of IPS seeds sharing the same mediolateral or anteroposterior position, and averaging connectivity profiles across seeds within each group (Fig. 3), thereby summarizing connectivity along each IPS spatial axis. This procedure yielded 11 mediolateral seed groups and 18 anteroposterior seed groups, corresponding to the discrete IPS positions represented in the seed layout. For the anteroposterior analysis, only the 11 mediolateral seed positions spanning m5 to l5, which were represented at all 18 anteroposterior levels, were included to ensure comparable sampling across AP positions. This reduction was used to facilitate visualization of the overall spatial relationship between IPS seed position and S1 connectivity, rather than to assume that connectivity was uniform across seeds at finer spatial scales. Connectivity profiles were then plotted as a function of S1 mediolateral position, allowing direct comparison across IPS seed groups (Fig. 3B).

**Figure 3.**
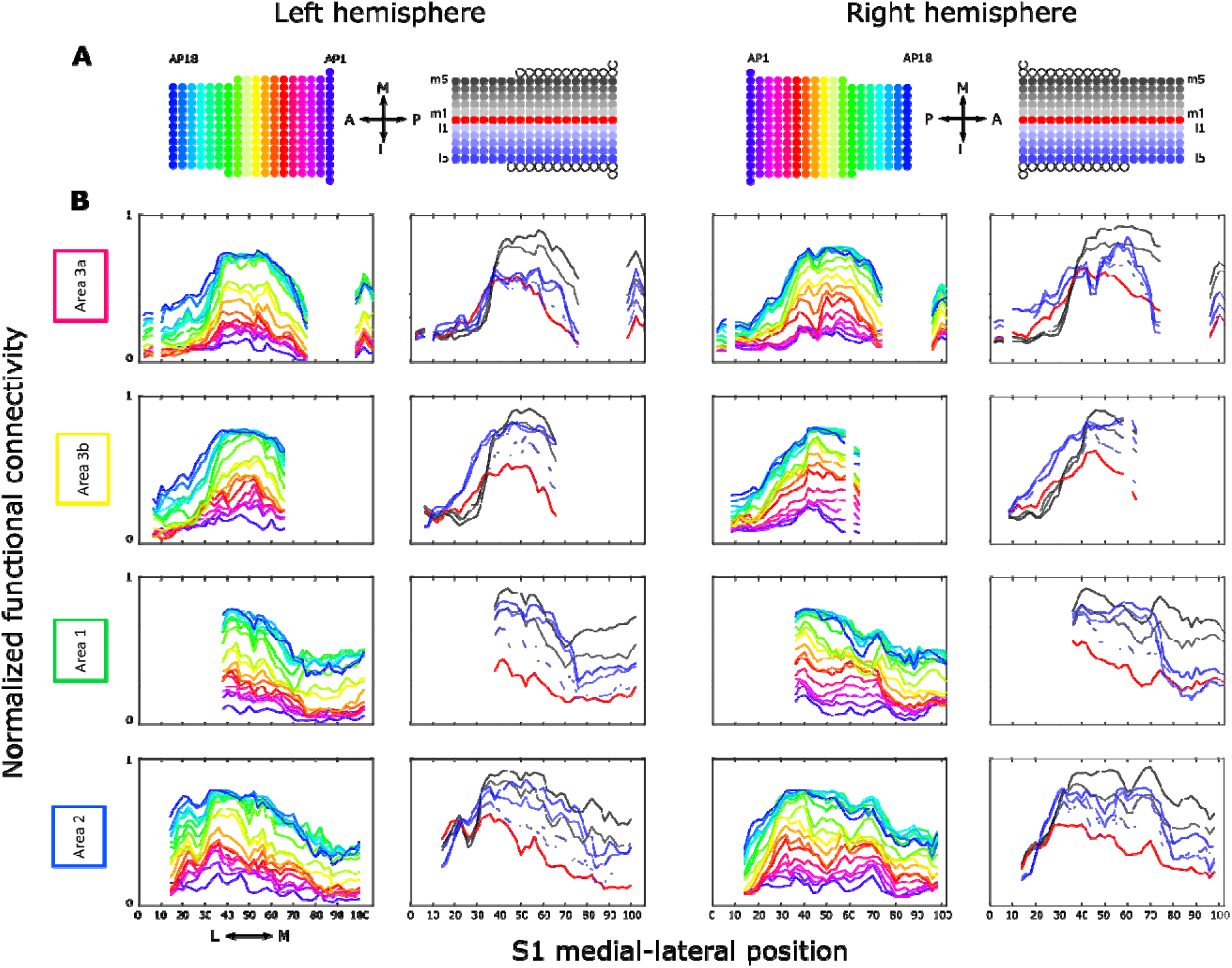
Systematic probing of positive IPS-S1 functional connectivity by varying IPS seed positions. **A.** Color legend for panel B, illustrating averaging positive functional connectivity patterns along either the anteroposterior axis (AP1-AP18; represented by different hues) or the mediolateral axis (medial positions m1-m5 shown in black, lateral positions l1-l5 shown in blue with varying values, and fundus positions shown in red) in both hemispheres. For the mediolateral analysis, posterior seeds that did not span the full anteroposterior extent of the IPS were excluded and are indicated by empty circles. A, anterior; P, posterior; M, medial; L, lateral. **B.** Normalized positive-connectivity profiles (y axis) along the two IPS spatial axes for areas 3a, 3b, 1, and 2. The x-axis represents the mediolateral positions of S1 on the cortical flatmap: 1 is lateral most S1 positions; 120 is medial most S1 positions.

Because IPS seeds located more anteriorly were also generally closer to S1, we performed an additional analysis to assess the contribution of anatomical proximity to the anteroposterior connectivity gradient. For each hemisphere and S1 subregion, three-dimensional Euclidean distance was calculated from each IPS seed to the vertices of the corresponding S1 subregion on the mid-thickness cortical surface. The primary distance measure was the mean Euclidean distance from each IPS seed to all vertices within that S1 subregion. For each of the 18 anteroposterior IPS positions, this distance was then averaged across the same 11 mediolateral seed positions (m5–l5) used in the Fig. 3 anteroposterior analysis.

For each monkey separately, the Fig. 3 anteroposterior connectivity profile was reconstructed using the same binarized positive-connectivity data and spatial averaging procedure as in the visualization. A single connectivity value was obtained for each AP position by averaging the resulting profile across S1 mediolateral positions. We then fitted two linear models within each monkey: an unadjusted model containing AP position as the predictor and a distance-adjusted model containing both AP position and mean Euclidean distance to S1. AP position and distance were standardized before model fitting. The AP regression coefficients obtained from the 10 monkeys were tested against zero using one-sample t-tests. To account for testing across four S1 subregions and two hemispheres, p values for the distance-adjusted AP coefficients were corrected using the Holm procedure. Spearman correlations between AP position and Euclidean distance, together with variance inflation factors (VIFs), were used to characterize collinearity between the two predictors. As sensitivity analyses, the models were repeated using the distance from each IPS seed to the nearest S1 vertex and the distance to the centroid of the corresponding S1 subregion instead of mean distance (Supplementary Fig. S1).

#### 3.3.3 Unsupervised clustering and robustness analyses

To further characterize similarities among IPS–S1 connectivity patterns without imposing predefined spatial constraints, we applied unsupervised hierarchical clustering independently within each S1 subregion (areas 3a, 3b, 1, and 2). Pairwise dissimilarity between IPS seeds was quantified using the Chebyshev distance, and clusters were constructed using average linkage. Because hierarchical clustering does not define a unique number of clusters, clustering resolution was determined using an iterative size-based criterion. Beginning with four clusters, k and (k+1)-cluster solutions were compared sequentially, and additional subdivisions were retained when the newly introduced groups exceeded 5% of all IPS seeds and were larger than the smallest group in the current solution. Iteration was otherwise terminated when the smallest group in the (k+1) solution was at or below 5%. To evaluate sensitivity to this criterion, the analysis was repeated using 10% and 2.5% thresholds while keeping all other clustering parameters unchanged (Supplementary Fig. S2). The 5% solution was used for the primary analysis and visualized using dendrograms, cluster-labeled IPS seed maps, and averaged connectivity profiles along the S1 mediolateral axis (Fig. 4). As a quality-control analysis, the partitions obtained using the dendrogram-based cutting procedure were compared with those obtained using MATLAB’s direct maximum-cluster criterion; the resulting partitions were identical in all S1 subregions and hemispheres (ARI = 1).

**Figure 4.**
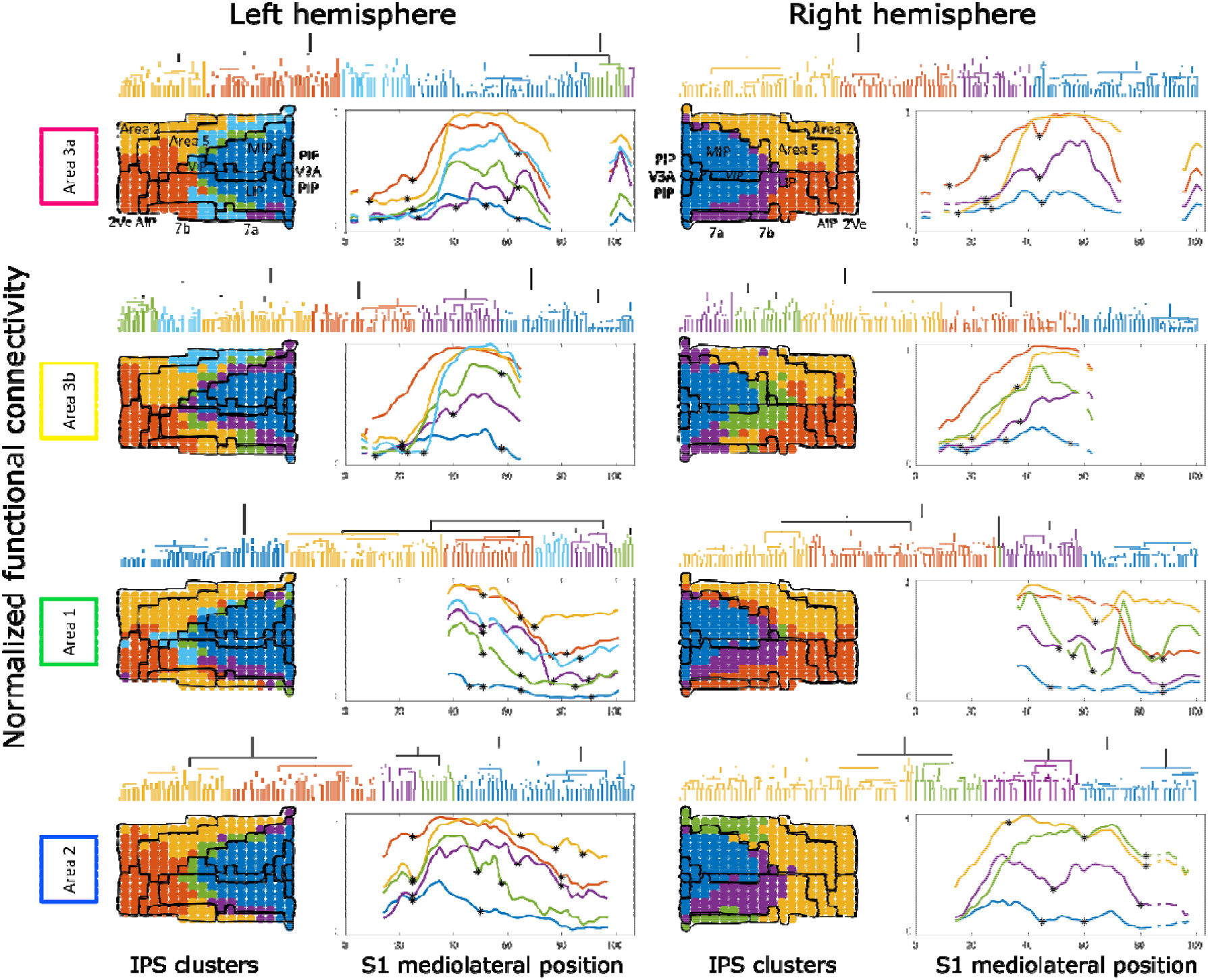
Unsupervised clustering of IPS-S1 positive functional connectivity patterns. In each S1 subregion (areas 3a, 3b, 1, and 2) of both hemispheres, the 223 IPS-S1 positive-connectivity profiles were grouped into four to six clusters using hierarchical clustering. Dendrograms illustrate the clustering results. IPS seeds are color-coded according to cluster membership and displayed relative to areal boundaries defined by the Markov atlas (Markov et al., 2014). Averaged normalized positive-connectivity profiles for each cluster are plotted as a function of S1 mediolateral position, with stars indicating local minima retained according to the criteria described in Methods 3.3. The x-axis represents the mediolateral positions of S1 on the cortical flatmap: 1 is the lateral most S1 position; 120 is the medial most S1 position.

Because the seed-placement strategy resulted in shorter inter-seed distances near the IPS fundus, we additionally assessed whether unequal spatial sampling influenced the connectivity-defined clustering. Three-dimensional Euclidean distances were calculated between IPS seed locations, and randomized spatial thinning was used to reduce the overrepresentation of densely sampled regions. For each hemisphere, 500 thinning realizations were generated using minimum inter-seed separation criteria of 1.5 and 2.0 mm. For each realization, hierarchical clustering was repeated independently within areas 3a, 3b, 1, and 2 using the same distance metric, linkage method, and iterative cluster-number procedure as in the primary analysis. The 5% minimum cluster-size criterion was calculated relative to the number of seeds retained in each thinning realization. Similarity between each recomputed partition and the corresponding original clustering, restricted to the same retained seeds, was quantified using the adjusted Rand index (ARI), which is invariant to arbitrary cluster labels and adjusted for chance agreement (Supplementary Fig. S3).

#### 3.3.4 Correspondence with cytoarchitectonic organization and individual-level boundary reproducibility

To assess the correspondence between connectivity-defined IPS clusters and cytoarchitectonic organization, each of the 223 IPS seed positions was assigned to a Markov atlas area based on visual comparison of the seed location with the atlas in volumetric space. These manually assigned area labels were digitized on the same unfolded IPS seed lattice used for visualization. Correspondence between the functional cluster partition and the resulting Markov atlas partition was quantified separately for each S1 subregion and hemisphere using adjusted mutual information (AMI) as the primary measure. Normalized mutual information (NMI) and adjusted Rand index (ARI) were additionally calculated as complementary measures of partition similarity.

We also calculated cluster-to-atlas purity, defined as the proportion of seeds within each functional cluster belonging to its most represented Markov area, averaged across clusters according to cluster size. AMI and ARI account for agreement expected by chance, whereas NMI quantifies the normalized information shared between the two partitions. Because both the functional clusters and atlas areas were spatially structured, no inferential test based on random permutation of individual seed labels was performed. Seeds without an unambiguous anatomical assignment were excluded from this analysis (Supplementary Fig. S4).

To assess whether the group-defined functional boundaries were consistently expressed in individual animals, hierarchical clustering was additionally performed separately for each monkey, S1 subregion, and hemisphere using the same Chebyshev distance and average-linkage procedure as in the group analysis. For this analysis, the number of clusters was fixed to the number identified in the corresponding group-level 5% solution, thereby assessing variation in boundary location while holding clustering resolution constant. Individual dendrograms were therefore cut using MATLAB’s maximum-cluster criterion with the corresponding group-level value of k.

Boundary reproducibility was quantified on the unfolded IPS seed lattice (Supplementary Fig. S5A). Pairs of cardinally adjacent seeds were enumerated, and a functional boundary edge was defined when the two adjacent seeds were assigned to different clusters. For each group-defined boundary edge and each monkey, we calculated the Euclidean distance between the midpoint of the group boundary edge and the midpoint of the nearest boundary edge in the corresponding individual-animal clustering solution. Distances were calculated from the anteroposterior seed position and mediolateral seed index and are therefore expressed in unfolded lattice units rather than millimeters, because physical inter-seed spacing was non-uniform across IPS. A distance of zero indicated an exact boundary-edge match, whereas progressively larger values indicated greater spatial separation between the group-defined and individual-animal boundaries.

To determine whether boundary reproducibility varied according to anatomical location, each group boundary edge was categorized according to the digitized Markov atlas labels of its two endpoint seeds. Edges whose endpoints belonged to the same atlas area were classified as within-area functional subdivisions, whereas edges joining two different atlas areas were classified as between-area functional boundaries. For each monkey and anatomical context, nearest-boundary distances were first summarized by taking the median across relevant boundary edges separately within each hemisphere × S1-subregion combination and then by taking the median across available combinations, yielding one value per monkey and anatomical context. Inferential analyses were restricted to anatomical contexts represented by at least five distinct group-defined boundary edges and for which values were available for all 10 monkeys. Within-area and between-area boundary contexts were treated as two predefined analysis families (Supplementary Fig. S5B-C). Differences among anatomical contexts within each family were first assessed using Friedman repeated-measures tests. The two omnibus p values were corrected using the Holm procedure. When the corresponding omnibus test was significant, pairwise comparisons were performed using Wilcoxon signed-rank tests, with Holm correction applied separately across all pairwise comparisons within each analysis family. For visualization, individual monkey values and group medians were plotted for the within-area and between-area analyses. Boundary reproducibility was additionally displayed separately for each hemisphere × S1-subregion combination as a heatmap, with each cell calculated by first taking the median across relevant boundary edges within each monkey and then the median across monkeys (Supplementary Fig. S5D). Anatomical contexts not represented by a group-defined boundary in a given hemisphere × S1-subregion combination were left unassigned in the heatmap.

#### 3.3.5 Identification of putative S1 somatotopic boundaries

The unsupervised clustering approach also allowed us to examine whether spatial transitions in IPS-S1 positive connectivity reflected somatotopic boundaries in S1 by identifying prominent local connectivity minima in smoothed connectivity profiles along the mediolateral axis. Candidate local minima were identified in the smoothed cluster-averaged connectivity profiles. For each local minimum, the differences in connectivity between the minimum and its adjacent local maxima were calculated. A minimum was retained as a putative boundary when either the peak-to-valley difference on the preceding or following side exceeded the mean peak-to-valley difference across the corresponding set of candidate transitions by 0.5 standard deviations (Fig. 4) and projected back onto the S1 flatmap for comparison with body-part boundaries reported in the literature (Figs. 5, S6).

**Figure 5.**
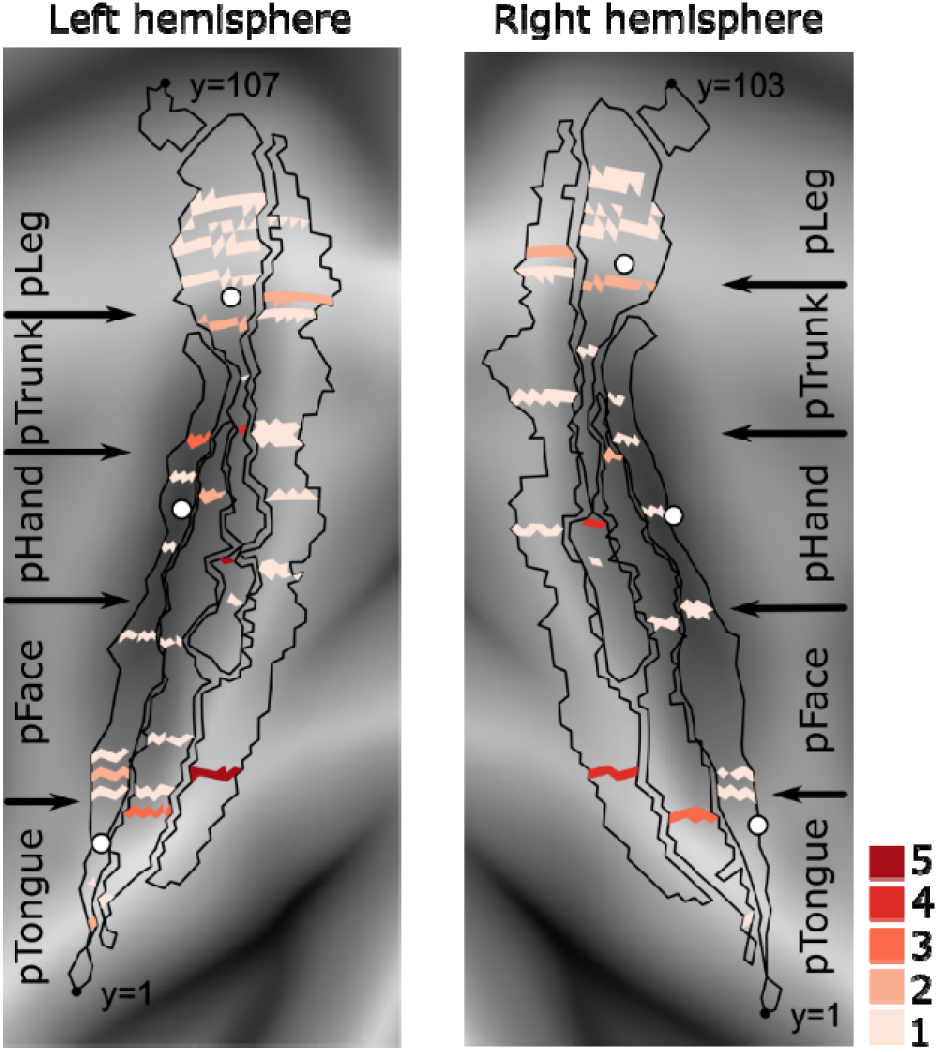
Local minima of IPS-S1 positive connectivity identify candidate somatotopic boundaries in S1. Local minima identified from connectivity profiles in areas 3a, 3b, 1, and 2 were retained according to the filtering criteria described in Methods 3.3. The mediolateral S1 positions of the selected local minima were projected onto the original cortical flatmaps together with the outlines of the S1 subregions in both hemispheres. The five-level red color scale indicates the multiplicity of local minima across distinct connectivity clusters at each mediolateral S1 position, with progressively darker shades representing 1, 2, 3, 4, or 5 connectivity clusters whose profiles contained a local minimum at that position. Thus, darker locations indicate positions at which local minima converged across a larger number of independent connectivity profiles from different IPS subregions. Putative somatotopic regions are labelled with a “p” prefix and include putative tongue, face, hand, trunk, and leg representations. Solid arrows indicate local minima that showed correspondence with putative somatotopic boundaries (also see Fig. S6).

#### 3.3.6 Somatotopic composition of IPS connectivity clusters

To further characterize the somatotopic composition of the IPS connectivity clusters, the cluster-averaged connectivity profiles were divided along the S1 mediolateral axis into five putative body-part intervals corresponding to tongue, face, hand, trunk, and leg representations. The boundaries between these intervals were defined separately for the two hemispheres based on the local minima corresponding to putative somatotopic transitions identified in the preceding analysis (Figs. 5 and S6). For each IPS cluster and each S1 subregion (areas 3a, 3b, 1, and 2), connectivity strength for a given body-part interval was quantified as the mean cluster connectivity across S1 positions within that interval and normalized by the number of monkeys. Mean rather than summed connectivity was used to prevent wider somatotopic intervals from receiving larger values solely because they contained more sampled positions.

When an IPS seed group exhibited significant connectivity with one of the S1 body part regions defined as described above, we interpreted this connectivity as evidence that the corresponding IPS region may contain a representation of that body part, and refer to it as the respective IPS body-part representation. For visualization (Fig. 6), these body-part representations were displayed as pictograms within each IPS cluster. We labeled as representation any seed group with a connectivity strength of at least 25% of the strongest body-part connectivity within the corresponding cluster and at least 8% of the maximum connectivity strength across all clusters and S1 subregions within the corresponding hemisphere, with a maximum of three pictograms displayed per cluster. When no representation satisfied these criteria, we labeled the seed group with the body-part that exhibited the strongest correlation. Pictogram area was scaled according to normalized connectivity strength. Pictograms were positioned within the largest spatially connected component of each cluster for visualization only; therefore, their precise position within a cluster does not indicate the cortical location of the corresponding body-part representation. IPS areal boundaries from the Markov atlas were overlaid on the cluster maps to facilitate comparison between functional clustering and anatomical organization.

**Figure 6.**
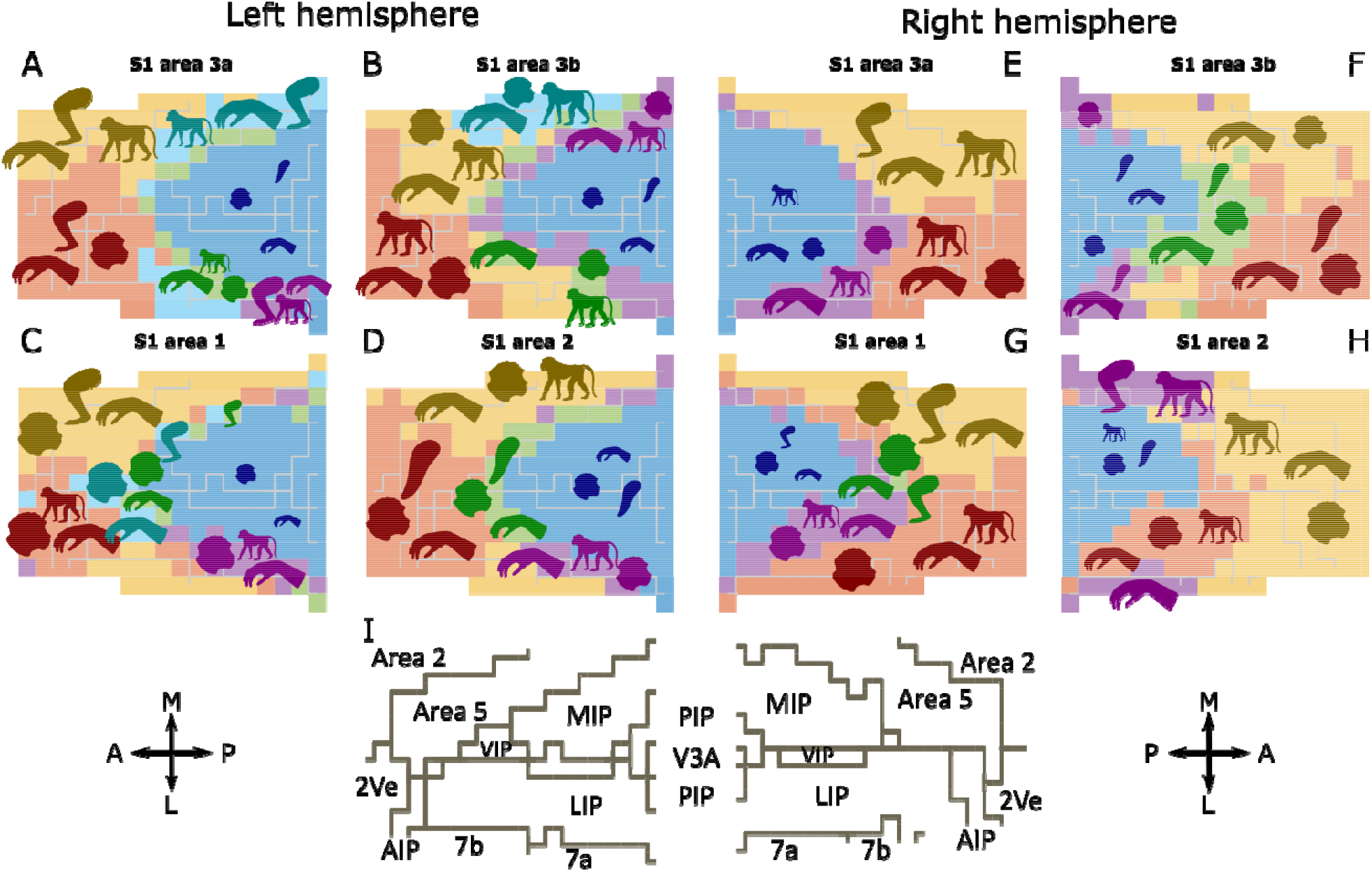
Somatotopic composition of IPS functional-connectivity clusters across S1 subregions. **A-D.** Spatial organization of IPS connectivity clusters derived from functional-connectivity profiles with left-hemisphere S1 areas 3a (A), 3b (B), 1 (C), and 2 (D), respectively. Colored patches indicate IPS cluster membership, and gray lines indicate anatomical areal boundaries within the IPS. Within each cluster, body-part pictograms summarize its strongest connectivity with putative S1 somatotopic regions. The S1 mediolateral axis was divided into five putative somatotopic intervals corresponding to tongue, face, hand, trunk, and leg representations. For each cluster and body-part interval, connectivity strength was quantified as the mean connectivity across S1 positions within that interval, normalized by the number of monkeys. Up to three body-part pictograms are displayed for each cluster. Pictograms were retained when their connectivity strength was at least 25% of the strongest body-part connectivity within that cluster and at least 8% of the maximum connectivity strength across the four S1 subregions; when no pictogram met these criteria, the strongest body-part representation was retained. Pictogram size reflects normalized connectivity strength, with larger pictograms indicating stronger connectivity. Pictogram color is derived from the corresponding cluster color for visualization and does not represent an additional quantitative variable. **E-H**. Same as in A–D, for right-hemisphere S1 areas 3a (E), 3b (F), 1 (G), and 2 (H), respectively. **I.** Anatomical reference maps showing the IPS areal boundaries used to interpret the functional clusters in the left and right hemispheres, based on the Markov atlas. Anatomical area labels correspond to the atlas-defined subdivisions of the IPS and surrounding parietal cortex. A, anterior; P, posterior; M, medial; L, lateral. AIP, anterior intraparietal area; LIP, lateral intraparietal area; MIP, medial intraparietal area; PIP, posterior intraparietal area; VIP, ventral intraparietal area.

### 3.4 Temporal signal-to-noise ratio assessment

Temporal signal-to-noise ratio (tSNR) was calculated voxel-wise for each monkey as the temporal mean signal divided by its temporal standard deviation. Because the functional data were already represented in F99 template space, individual-monkey tSNR maps were averaged to obtain a group-level tSNR map. For whole-brain surface visualization, the group-average volumetric tSNR map was projected onto the cortical surface using Connectome Workbench with ribbon-constrained volume-to-surface mapping based on the corresponding white, pial, and mid-thickness surfaces (Marcus et al., 2013) (Fig. S7A-C).

To assess signal quality specifically at the IPS locations used for the functional connectivity analysis, tSNR was additionally sampled at the F99-space locations corresponding to the 223 single-voxel IPS seeds in each hemisphere. The corresponding tSNR value was extracted for each seed and each monkey from the individual-monkey volumetric tSNR maps using nearest-neighbor sampling to avoid introducing additional interpolation, and seed-wise values were averaged across monkeys and visualized using the same unfolded 223-seed IPS spatial organization used for the functional connectivity analyses. For quantitative comparison across the sulcus, tSNR values were averaged within medial, fundal, and lateral IPS seed groups for each monkey. Fundal tSNR was compared with medial and lateral IPS tSNR using paired two-sided t-tests across monkeys, with Holm correction for the two comparisons performed within each hemisphere. Because seed-level tSNR values were extracted directly from the volumetric tSNR maps, these quantitative analyses did not depend on the volume-to-surface projection used for visualization.

We further tested whether local signal quality was generally related to IPS-S1 functional connectivity across the full set of IPS seed positions. For each monkey and hemisphere separately, local tSNR at each of the 223 IPS seeds was correlated with the corresponding seed’s mean functional connectivity to S1. Mean connectivity was calculated across all structurally valid mediolateral positions within S1 areas 3a, 3b, 1, and 2. Spearman rank correlation was used as the primary measure because no linear relationship between tSNR and connectivity was assumed. This yielded one correlation coefficient per monkey and hemisphere. The resulting coefficients from the 10 monkeys were tested against zero using two-sided Wilcoxon signed-rank tests, thereby treating monkeys rather than individual IPS seed locations as the unit of statistical inference. P values were corrected across the two hemispheres using the Holm procedure (Fig. S7D-F). As a complementary analysis consistent with the binarized connectivity representation used in the main IPS-space analyses, the same procedure was repeated after expressing connectivity for each IPS seed as the proportion of structurally valid S1 mediolateral positions showing significant positive connectivity (Fig. S7G).

## Results

In primates, the anterior parietal somatosensory cortex, often collectively referred to as S1, comprises areas 3a, 3b, 1, and 2 (Kaas et al., 1979; Delhaye et al., 2018). These areas differ in their dominant sensory inputs and response properties. Area 3a is dominated by proprioceptive signals, including input related to muscle stretch and joint movement; area 3b is predominantly cutaneous and supports high-resolution spatial processing of tactile stimuli; area 1 remains primarily cutaneous and is particularly implicated in the processing of texture and surface features; and area 2 contains convergent cutaneous and proprioceptive representations important for encoding hand configuration, object size, and object shape (Carlson, 1981; DiCarlo et al., 1998; Delhaye et al., 2018). Area 2 also has direct corticocortical connections with the medial and lateral subdivisions of posterior parietal area 5, areas 5M and 5L, respectively (Padberg et al., 2019). This progression from predominantly proprioceptive processing through cutaneous feature processing to multimodal integration provides a useful framework for interpreting differential IPS-S1 functional connectivity across S1 subregions.

To characterize this differential IPS-S1 functional connectivity across S1 subregions at a fine spatial scale, we analyzed awake resting-state fMRI data from 10 macaque monkeys. A total of 223 single-voxel seeds were placed throughout the IPS, including areas 2, 2Ve, 5, 7a, 7b, AIP, VIP, LIP, MIP, PIP, and V3A, across both the medial and lateral banks in the two hemispheres (Fig. 1A-D, F). Seed-to-whole-brain functional connectivity was calculated in volumetric space and then projected onto the cortical surface (see Methods 3.3). We used the Paxinos atlas (Paxinos et al., 1999) to identify the four S1 subregions, area 3a, 3b, 1 and 2 (Fig. 1E). In the following, we focused on functional connectivity between IPS and S1. Functional connectivity refers to the binarized presence of significant positive connectivity unless otherwise specified.

### Mediolateral position within S1 shapes the topography of IPS functional connectivity

We first examined how IPS-S1 functional connectivity varies as a function of the mediolateral position along S1. To this end, we applied a fixed-size sliding window (Fig. 2C) to each S1 subregion (areas 1, 2, 3a and 3b) and quantified the spatial distribution of functional connectivity between each window and each IPS voxel (Fig. 2A, B, D). The four S1 subregions showed both shared and distinct patterns of IPS connectivity as the sliding window progressed from lateral to medial positions.

**Area 3a** (proprioceptive input from muscle spindle afferents). At the most lateral positions, functional connectivity was generally sparse across the IPS, with no region exhibiting substantially elevated connectivity prevalence. As the window shifted medially, a semi-oval region of elevated connectivity emerged, surrounding a region of relatively lower connectivity within the IPS. Connectivity was initially elevated at the anterior tip of the IPS and progressively extended along its perimeter, while connectivity remained reduced near the fundus except at the anterior tip. The semi-oval appeared approximately four successive sliding-window positions further medially than in area 2. Within this pattern, connectivity was slightly more pronounced along the medial IPS bank as compared to its lateral bank. With further medial shifts, the semi-oval region of elevated connectivity gradually became less distinct and gradually dissolved into a single region of strong connectivity confined to the medial IPS bank, as the low-connectivity zone expanded preferentially toward the lateral bank.

**Area 3b** (cutaneous inputs encoding fine spatial detail). The most lateral positions showed enhanced connectivity in the lateral bank of the anterior IPS. More medially, a symmetric pattern along the perimeter of IPS emerged, as observed in area 3a. The semi-oval structure of elevated connectivity showed the highest functional connectivity among areas 3a, 1 and 2, up to the most medial area 3b positions.

**Area 1** (cutaneous information related to texture and object properties). Among the four subregions, area 1 showed the most lateral onset of enhanced anterior IPS connectivity: it was already evident in the most lateral sliding window, with connectivity prevalence appearing qualitatively higher there than at the more medial sliding-window positions within area 1. As the window shifted medially, the anteroposteriorly elongated region of low connectivity expanded preferentially toward the lateral bank, leaving a single region of strong connectivity in the medial IPS, a progression similar to area 3a and occurring at the same mediolateral axis.

**Area 2** (combining proprioceptive and cutaneous signals). A strong functional connectivity was observed between the most lateral area 2 sliding-window position and the anterior tip of the IPS. As the sliding window shifted medially, the location of strong connectivity shifted slightly toward the lateral end of the IPS. For more medial area 2 sliding-window positions, a semi-oval region of elevated connectivity appeared at relatively lateral positions, whereas a similar pattern emerged at more medial sliding-window positions in areas 3a, 3b, and 1. As the window shifted medially, the transition out of the semi-oval pattern also appeared to occur at a more lateral sliding-window position than in areas 1 and 3b. The anteroposteriorly elongated low-connectivity region expanded relatively symmetrically across the medial and lateral banks, distinguishing area 2 from areas 1 and 3a, where the corresponding regions expanded asymmetrically. Toward the most medial area 2 sliding-window positions, connectivity became more pronounced along the medial convexity of the IPS.

In summary, IPS-S1 functional connectivity varied systematically as a function of the mediolateral position within S1. A common feature observed across all S1 subregions was a semi-oval connectivity pattern, characterized by elevated prevalence of strong connectivity along the IPS perimeter and reduced connectivity near the fundal IPS, except at the anterior tip. This semi-oval pattern closely followed the geometry of the intraparietal sulcus itself. Beyond this shared organization, qualitative differences were observed across S1 subregions in the mediolateral position at which the semi-oval emerged, in the relative contributions of the medial and lateral IPS banks, and in the persistence or dissolution of the pattern at medial positions. Together, these observations suggest subregion-specific organization of IPS-S1 connectivity, which is examined further in the analyses below.

### IPS-S1 connectivity follows systematic anteroposterior and mediolateral gradients across all S1 subregions

The sliding-window analysis characterized IPS-S1 connectivity from the perspective of S1, revealing how the spatial distribution of IPS connectivity changes as a function of mediolateral position within each S1 subregion. We next examined the same connectivity patterns from the complementary perspective of IPS, asking how S1 connectivity varies as a function of IPS seed position along the anteroposterior and mediolateral axes. To this end, we averaged connectivity profiles across seeds sharing the same anteroposterior or mediolateral position within the IPS (Fig. 3), exploiting the fact that S1 dimensionality had already been reduced to a single mediolateral axis (see Methods 3.3). This averaging provides a reduced representation of the connectivity patterns; although useful for revealing broad spatial gradients, it may obscure finer-grained variations among seeds at the same anteroposterior or mediolateral position.

Following this strategy, we first averaged functional connectivity according to the anteroposterior position of IPS seeds and normalized to the maximum connectivity observed for each S1 subregion (Fig. 3, first and third columns). Across all four S1 subregions, a clear ordering of normalized connectivity was observed, along the anatomical anteroposterior axis of the IPS: functional connectivity was greatest for anterior IPS seeds and progressively decreased toward posterior IPS seeds. In addition, all subregions exhibited a broad plateau of elevated connectivity centered along the middle portion of the S1 mediolateral axis. Although areas 3a, 3b, 1, and 2 occupy distinct mediolateral positions within S1, this plateau-like pattern was consistently present across all subregions. Nevertheless, subtle differences were also observed across subregions. For example, the plateau was broader and rose more gradually from lateral positions in areas 2 than in areas 3a and 3b.

Because anterior IPS positions were also generally closer to S1, we next examined whether this anteroposterior connectivity gradient could be accounted for by anatomical proximity (Fig. S1). Using the same IPS positions included in the Fig. 3 anteroposterior analysis, we estimated the AP effect separately for each monkey before and after including mean Euclidean distance to the corresponding S1 subregion as a covariate. After correction for multiple comparisons, the AP effect remained significant in area 1 bilaterally and in left-hemisphere area 2. In areas 3a and 3b, however, AP position and Euclidean distance were highly collinear, limiting the ability to statistically distinguish their independent contributions. These results indicate that anatomical proximity contributes to the observed anteroposterior organization but does not fully account for it, particularly in area 1.

We then averaged functional connectivity according to the mediolateral position of IPS seeds (Fig. 3, second and fourth columns). A similarly ordered pattern of normalized connectivity emerged. Connectivity associated with fundus seeds (red) was generally weakest, with the exception of in the lateral portion of S1. In contrast, connectivity patterns from the medial (black) and lateral (blue) IPS banks were largely symmetric: connectivity decreased toward the fundus and increased toward the cortical convexities. In all areas, the most medial IPS showed higher connectivity to all S1 areas than the most lateral IPS. This was particularly the case for area 3a. Consistent with the previous analysis, the plateau of elevated connectivity along the middle portion of the S1 mediolateral axis was also clearly preserved.

The connectivity pattern near the lateral portion of S1 described above was notable. In this region, particularly within areas 3a and 3b, connectivity patterns from the medial and lateral IPS banks were asymmetric. Specifically, functional connectivity associated with medial-bank seeds was even lower than that of the fundus seeds, whereas connectivity from lateral-bank seeds remained the strongest. However, as S1 positions shifted medially, connectivity associated with medial-bank seeds increased more steeply toward the central plateau and eventually exceeded that of the lateral-bank seeds within the plateau region.

Taken together, averaging functional connectivity across IPS seeds along either the anteroposterior or mediolateral axis provided an effective dimensionality-reduction approach and enabled a systematic examination of IPS-S1 positional effects. The systematic increase in functional connectivity toward anterior IPS positions, together with elevated connectivity near the cortical convexities, was consistent with the semi-oval connectivity structure observed in the sliding-window analysis (Fig. 2). Likewise, the asymmetry between medial- and lateral-bank IPS seeds was consistent with the asymmetric expansion of the anteroposteriorly elongated low-connectivity region and the persistence of a single region of elevated connectivity within the medial IPS.

Because reduced functional connectivity near the IPS fundus could potentially reflect poorer local fMRI signal quality, we additionally examined temporal signal-to-noise ratio (tSNR; Fig. S7). Whole-brain group-average tSNR maps characterized the spatial distribution of signal quality across the cortex, and tSNR was further sampled at the 223 IPS seed locations in each hemisphere. We first tested whether local tSNR was generally associated with IPS–S1 connectivity across seed positions. For each monkey and hemisphere, Spearman correlations were calculated across the 223 IPS seeds between local tSNR and mean connectivity to S1 (Fig. S7). Although the strength and direction of these correlations varied across monkeys, they were not systematically different from zero in either hemisphere (left: median ρ = 0.25, Wilcoxon signed-rank p = 0.11; right: median ρ = −0.01, p = 0.77). A complementary analysis based on the prevalence of positive S1 connectivity yielded the same conclusion (left: median ρ = 0.14, p = 0.13; right: median ρ = 0.03, p = 0.77). We next examined tSNR specifically across the unfolded IPS organization. The reduced connectivity near the fundus was not accompanied by a corresponding reduction in local tSNR. Fundal tSNR was comparable to medial IPS and higher than lateral IPS in the left hemisphere, whereas it was higher than medial IPS and comparable to lateral IPS in the right hemisphere. Thus, neither the seed-wise relationship between tSNR and IPS–S1 connectivity nor the fundus-specific tSNR distribution supported poorer local signal quality as an explanation for the reduced fundal connectivity.

### Unsupervised clustering reveals a large-scale functional topology of IPS-S1 connectivity that only partially corresponds to cytoarchitectonic boundaries

The sliding-window analysis (Fig. 2) characterized IPS-S1 connectivity from the perspective of S1, revealing how the spatial distribution of IPS connectivity varies as a function of mediolateral position within each S1 subregion. The positional averaging analysis (Fig. 3) addressed IPS-S1 connectivity by probing IPS seed space along its two principal anatomical axes, revealing clear anteroposterior and mediolateral gradients. Yet, both approaches imposed predefined spatial decompositions that may not capture the full organizational structure of IPS-S1 connectivity. To characterize this structure without predefined assumptions, we next applied unsupervised hierarchical clustering directly to all 223 IPS-S1 positive-connectivity profiles (see Methods 3.3). This approach groups seeds by the similarity of their connectivity fingerprints, irrespective of their spatial position, allowing the data to reveal its own organizational structure. The resulting clusters were visualized using dendrograms, two-dimensional IPS seed maps, and normalized functional connectivity profiles plotted as a function of S1 mediolateral position (Fig. 4).

Using hierarchical clustering, with the 5% minimum cluster-size criterion used for the primary analysis, we identified four to six clusters in all S1 subregions and in both hemispheres. Across all subregions, the clustering consistently recovered the broad semi-oval IPS organization previously described in the sliding-window analysis (Fig. 2). In general, the clustering patterns comprised a central semi-oval sulcal cluster (dark blue), together with anterior-medial (yellow) and anterior-lateral (red) convexity clusters. Between the sulcal and convexity clusters, one to three intermediate nested transition bands were observed (purple, green, and cyan). The innermost purple band was located primarily along the posterior portions of the medial and lateral IPS banks, followed by a second green band near the anterior tip of the sulcus and, in some S1 subregions, a third cyan band positioned further outward. These intermediate clusters therefore occupied progressively more peripheral positions around the central semi-oval sulcal territory, partitioning the transition between sulcal and convexity connectivity patterns rather than defining sharply separated anatomical compartments. When the clustering was repeated using alternative minimum cluster-size criteria of 10% and 2.5%, the broad sulcal-to-convexity organization was preserved, whereas the number and spatial extent of the intermediate transition bands varied with clustering resolution (Fig. S2). Overall, the unsupervised analysis independently recovered the large-scale semi-oval organization identified by the preceding spatial analyses, while further subdividing the connectivity transition between its sulcal and convexity components.

Because IPS seeds were sampled more densely near the sulcal fundus than near the cortical convexities, we next assessed whether this non-uniform spatial sampling substantially influenced the clustering results (Fig. S3). Randomized spatial thinning using minimum inter-seed separations of 1.5 and 2.0 mm retained a median of 81 and 71 of 223 seeds in the left hemisphere, respectively, and 84 and 70 seeds in the right hemisphere. Despite this substantial reduction in sampling density, the recomputed cluster partitions remained substantially similar to the corresponding original solutions, with median adjusted Rand indices (ARIs) ranging from approximately 0.61 to 0.78 across S1 subregions and hemispheres. Similar levels of partition stability were observed for the 1.5- and 2.0-mm thinning criteria. In contrast, the exact number of clusters varied more across thinning realizations, indicating that the large-scale connectivity-defined organization was more robust to differences in seed sampling density than the finer level of subdivision. Together with the alternative cluster-size analysis, these results indicate that the broad semi-oval topology was robust, whereas the precise number of intermediate subdivisions was more dependent on clustering resolution and sampling configuration.

After identifying the hierarchical clusters, we examined how these clusters corresponded to cytoarchitectonic areal boundaries within IPS. Quantitative comparison with the Markov atlas revealed consistent but incomplete correspondence between the functional and cytoarchitectonic partitions (Fig. S4). Adjusted mutual information (AMI) ranged from 0.30 to 0.34 in the left hemisphere and from 0.32 to 0.43 in the right hemisphere. Complementary measures showed a similar pattern (NMI = 0.34–0.46; ARI = 0.16–0.33), indicating that the functional organization shared structure with the cytoarchitectonic parcellation but did not reproduce it one-to-one. Consistent with this partial correspondence, individual functional clusters commonly extended across multiple Markov atlas areas. The sulcal clusters (Fig. 4, dark blue) consistently encompassed areas MIP, VIP, V3A, PIP, and the posterior-medial portion of LIP across all of areas 3a, 3b, 1 and 2. The anterior-medial convexity clusters (Fig. 4, yellow) covered most of areas 5 and 2, but in several cases also extended into lateral IPS regions within LIP, 7a, and 7b (particularly in areas 3b, 1, and 2 of the left hemisphere). In contrast, the anterior-lateral convexity clusters (Fig. 4, red) included areas 2v, AIP, the anterior portion of LIP, and parts of area 7a, together with smaller lateral portions of areas 2 and 5. Thus, although connectivity-defined clusters showed systematic correspondence with established cytoarchitectonic organization, their boundaries frequently crossed individual atlas-defined areas.

The intermediate edge-layer clusters showed more selective correspondence with cytoarchitectonic boundaries. Most notably, they were frequently positioned near the boundary between MIP and area 5, while simultaneously subdividing LIP into posterior-medial and anterior-lateral compartments. This subdivision was consistently observed across S1 subregions and hemispheres. Although this group-level subdivision was observed across S1 subregions and hemispheres, its reproducibility in individual animals was spatially heterogeneous. To assess this directly, we compared each group-defined functional boundary with the nearest boundary obtained from fixed-k clustering of individual-animal connectivity profiles (Fig. S5). Boundary reproducibility differed significantly across anatomical contexts, both for functional subdivisions located within individual atlas areas and for functional boundaries spanning pairs of atlas areas (Friedman tests, both Holm-corrected p < 0.001). Within-area subdivisions in LIP were more consistently recovered across animals than those within VIP, area 5, and area 2 (all Holm-corrected p = 0.041), whereas the difference between LIP and MIP was not significant. Among between-area transitions, the 7b–LIP boundary was also more consistently preserved than the MIP–area 5 boundary (Holm-corrected p = 0.041), whereas the corresponding comparison between 7a–LIP and MIP–area 5 was not significant. These results indicate that the group-defined topology was not equally stable across IPS: boundaries involving the lateral convexity and LIP were generally more reproducible across individuals, whereas several medial and intermediate transitions, including the MIP–area 5 region, showed greater variability in their individual-level location. Because the cytoarchitectonic parcellation was defined from a common atlas rather than subject-specific histology, the apparent correspondence with individual areal boundaries should be interpreted as approximate. The observed variability in functional boundary location is distinct from possible inter-individual variability in the true cytoarchitectonic borders themselves, which cannot be directly assessed with the present data. Variation in true cytoarchitectonic borders across individuals, together with registration uncertainty, may therefore contribute to the observed mismatch between functional and atlas-defined boundaries. In addition, the intermediate clusters frequently included a small anterior portion of VIP in the left hemisphere, consistent with potential functional differentiation within this region as suggested by Sheng et al., 2025. Overall, these results indicate that IPS-S1 connectivity is organized according to large-scale functional topologies that only partially correspond to classical cytoarchitectonic areal boundaries.

### Local minima of IPS-S1 connectivity are associated with somatotopic transitions

Traditionally, somatotopic boundaries in S1 have been defined using electrophysiological recordings (Kaas et al., 1979; Nelson et al., 1980; Pons et al., 1985) and, more recently, functional imaging (Arcaro et al., 2019). Functional connectivity provides a complementary perspective on this organization by characterizing the intrinsic network architecture of the cortex. Whereas electrophysiological recordings and functional localizers identify body-part representations based on sensory responses, connectivity analyses reveal transitions in large-scale functional relationships. If neighboring somatotopic territories participate in partially distinct cortical networks, changes in connectivity may occur near representational transitions (Thomas et al., 2021) as described in M1 (Gordon et al., 2023). We therefore asked whether local reductions in IPS-S1 connectivity were preferentially located near atlas-defined somatotopic boundaries. Because these boundaries were derived from an independent anatomical/functional reference rather than from the same animals, such correspondence should be interpreted as approximate and may also reflect inter-individual variability in the location of representational boundaries.

The systematic variation of IPS-S1 functional connectivity revealed by the unsupervised hierarchical clustering analysis provided an opportunity to test this prediction. We first identified local minima in the cluster-averaged connectivity profiles and retained the most prominent minima according to the criteria described in Methods 3.3 (stars in Fig. 4). The mediolateral S1 positions of these minima were projected onto the cortical flatmap for each S1 subregion (Fig. 5) and compared with previously published functional localizers (Fig. S6, Arcaro et al. (2019), Wardak et al. (2016), Gordon et al., 2023, Thomas et al., 2021). Several minima corresponded closely to reported transitions between putative tongue (Bono et al., 2022), face, hand/trunk, and leg representations (Nelson et al., 1980; Sur et al., 1982; Jain et al., 1997).

The clearest correspondences were observed at the putative hand/trunk-leg boundary in areas 1 and 2, and at the tongue-face boundary in areas 3a, 3b, and 1. In contrast, the face-hand boundary, which has attracted considerable attention in studies of S1 organization and plasticity, was less clearly reflected by the connectivity-derived minima. Because the face-hand boundary has been particularly prominent in studies of S1 organization and plasticity (Manger et al., 1997), the tongue-face boundaries identified in areas 3a, 3b, and 1 were initially considered as possible candidates for the face-hand transition. However, their mediolateral positions were substantially more lateral than expected based on both published functional localizers (Fig. S6) and classical electrophysiological maps (Kaas et al., 1979; Nelson et al., 1980; Delhaye et al., 2018), suggesting that these minima are more likely associated with the tongue-face boundary than with the face-hand boundary. Only a local minimum in area 3b of the right hemisphere corresponded closely to the published face-hand boundary (Fig. 5), whereas the corresponding minima in areas 1 and 2 of the left hemisphere were located further medially.

Although not every reported somatotopic boundary was recovered consistently across S1 subregions and hemispheres, the prominent connectivity-derived minima showed spatial correspondence with somatotopic boundaries identified by electrophysiological recordings and functional localizers (Fig. S6). Together, these results indicate that IPS-S1 connectivity minima can capture spatial transitions corresponding to specific somatotopic boundaries, most consistently the hand/trunk-leg and tongue-face transitions.

### Heterogeneous somatosensory input contributes to functional diversity among IPS-S1 connectivity clusters

Having identified candidate somatotopic boundaries based on the spatial correspondence between connectivity-derived local minima and independently reported S1 somatotopic maps, we next used these boundaries to characterize the somatotopic composition of each connectivity-defined IPS cluster (Fig. 6; see Methods 3.3). For each S1 subregion, the cluster-averaged connectivity profile was divided into putative tongue, face, hand, trunk, and leg intervals, and the mean connectivity within each interval was quantified. Body-part pictograms in Fig. 6 summarize the strongest somatotopic contributions to each IPS cluster, with pictogram size reflecting connectivity strength. The resulting maps showed that the connectivity-defined IPS clusters were generally not associated with a single body-part representation. Instead, individual clusters exhibited connectivity with multiple somatotopic sectors, and the relative contribution of these sectors varied across S1 areas 3a, 3b, 1, and 2 and between clusters. At the same time, clusters occupying similar portions of the IPS frequently retained related somatotopic profiles across S1 subregions, indicating that the large-scale cluster organization identified in Fig. 4 coexists with variation in the body-part composition of their S1 connectivity. Thus, the functional diversity among IPS-S1 connectivity clusters reflects not only differences in their spatial position within IPS but also differences in the somatotopic composition of their S1 inputs (Fig. 6).

## Discussion

In the present study, we used dense resting-state fMRI sampling across an IPS-centered region of posterior parietal cortex to characterize its functional connectivity with the four subregions of primary somatosensory cortex (S1). Three principal findings emerged. First, IPS-S1 connectivity exhibited a highly structured spatial organization, including a prominent semi-oval topology that closely followed the geometry of the intraparietal sulcus and systematic gradients along both the anteroposterior and mediolateral dimensions of IPS. Second, unsupervised clustering revealed connectivity-defined subdivisions that only partially corresponded to classical cytoarchitectonic boundaries, highlighting large-scale functional organization that extends across established cortical areas while also revealing heterogeneous connectivity patterns within individual regions such as LIP, suggesting functional specialization at a finer spatial scale. This observation raises the possibility that functional organization in IPS extends below the scale of conventional cortical areas, consistent with recent evidence for spatially clustered mesoscale functional units within macaque cortical areas (Zhu et al., 2025). Third, local minima in IPS-S1 connectivity profiles— that is, relative troughs in connectivity along the mediolateral S1 dimension—identified candidate areal transitions corresponding to several previously described somatotopic boundaries within S1. In sum, several features of the IPS-S1 connectivity profiles corresponded with cortical characteristics previously established using anatomical, electrophysiological, and functional imaging methods, providing convergent support for the biological relevance of the connectivity organization identified here. At the same time, connectivity profiles revealed systematic patterns of large-scale cortical organization that are not readily captured by traditional areal parcellations or electrophysiological maps alone, consistent with the broader use of connectivity profiles to identify functionally distinct cortical organization (Johansen-Berg et al., 2004). Together, these findings suggest that resting-state functional connectivity provides a complementary perspective on the functional relationship between IPS and S1, capturing organizational features across multiple spatial scales, from connectivity patterns extending across established cortical areas to finer-scale heterogeneity within individual areas and transitions associated with somatotopic architecture.

The four cytoarchitectonically distinct subregions of S1 occupy a functional hierarchy that mirrors the complexity of somatosensory processing (Kaas et al., 1979). Area 3a, buried in the fundus of the central sulcus, is the primary cortical target of proprioceptive signals from muscle spindle afferents, relayed via the shell region of the caudal division of the ventral posterior lateral nucleus (VPLc) of the thalamus (Friedman and Jones, 1981; Jones, 1983). Area 3b, by contrast, receives dense thalamic input from the cutaneous “core” of VPLc and is considered the primary cortical receiving area for tactile signals, with the highest proportion of cutaneous-responsive cells, the smallest receptive fields, and the most complete somatotopic representation among S1 subfields (Kaas et al., 1979). Areas 1 and 2 sit further from primary thalamic input and receive much of their driving input via feedforward projections from areas 3a and 3b, rather than from thalamus directly (Pons and Kaas, 1986; Garraghty et al., 1990). Neurons in area 1 respond almost exclusively to cutaneous stimulation, while neurons in area 2 respond to both cutaneous stimulation and stimulation of deep receptors of the skin and joints, consistent with its role in multimodal somatosensory integration and stereognosis (Yau et al., 2013). Area 2 occupies a pivotal position at the S1-parietal interface: its more restricted pattern of cortical connections indicates that it processes somatic inputs locally and provides proprioceptive information to area 5, which in turn is broadly connected with motor and posterior parietal areas (Padberg et al., 2019). Direct reciprocal connections between area 2 and IPS areas, including area 5 and, to a lesser extent, AIP and VIP, make it the most likely gateway through which somatosensory signals reach the IPS for sensorimotor integration. The relatively broader functional connectivity we observed between IPS and area 2 compared to areas 3a and 3b is therefore consistent with this anatomical hierarchy of S1–parietal projections. More broadly, the correspondence between the present functional connectivity patterns and known anatomical connectivity provides evidence that resting-state fMRI captures, at least in part, the signature of established cortico-cortical projection pathways, while also potentially reflecting interactions that extend beyond direct anatomical connections.

### Sulcus geometry is the prominent organizational feature of IPS-S1 connectivity

A principal finding of the present study is that IPS-S1 functional connectivity is organized according to a striking semi-oval topology, with relatively weak S1 connectivity within a central IPS territory surrounded by stronger connectivity, that closely follows the geometry of the intraparietal sulcus (Fig. 2 and 4). This organizational pattern emerged consistently across all four S1 subregions and in both hemispheres, indicating that it represents a robust feature of IPS-S1 interactions. Importantly, although all analyses were derived from the same underlying IPS-S1 connectivity dataset, the semi-oval topology was preserved across several complementary representations of the data, including sliding-window mapping (Fig. 2), positional averaging (Fig. 3), and unsupervised hierarchical clustering (Fig. 4). Such consistency suggests that the observed organization reflects a genuine property of the connectivity architecture rather than an artifact introduced by a particular visualization or dimensionality-reduction procedure.

The hierarchical clustering analysis (Fig. 4) provides particularly compelling support for this interpretation. Clustering was performed solely on the similarity of IPS-S1 connectivity profiles and did not incorporate any information about the anatomical positions of IPS seeds. Nevertheless, when the resulting cluster assignments were projected back onto IPS space, they reassembled into the same semi-oval arrangement revealed by the sliding-window analysis (Fig. 2). Thus, the semi-oval structure emerged directly from the connectivity relationships among IPS seeds rather than from their spatial arrangement. The broad sulcal-to-convexity organization was also preserved across alternative clustering resolutions and after substantial spatial thinning of the IPS seed set (Figs. S2 and S3), indicating that this large-scale topology was not strongly dependent on the precise cluster-size criterion or the non-uniform spatial density of the seed sampling. In contrast, the number and extent of finer intermediate subdivisions were more sensitive to these analytical choices.

The semi-oval topology further reflects systematic differences between IPS regions located near the cortical convexities and those buried within the sulcal fundus. The strongest positive connectivity with S1 was observed along the medial and lateral banks and convexities of the IPS, whereas connectivity was weakest near the fundus. Importantly, this reduction in fundal connectivity was not accompanied by a corresponding reduction in local temporal signal-to-noise ratio (tSNR; Fig. S7). Fundal tSNR was not lower than that of the surrounding medial and lateral IPS regions, arguing against poorer local fMRI signal quality as a trivial, non-neural explanation for the low fundal connectivity. In both the positional averaging and clustering analyses, the fundus region was consistently distinguished from the surrounding bank and convexity regions. Notably, the regions exhibiting the strongest S1 coupling were concentrated primarily within the anterior IPS, whereas connectivity generally weakened toward more posterior portions of the sulcus. Anatomical proximity to S1 contributed to this anteroposterior organization, although it did not fully account for the gradient, particularly in area 1 (Fig. S1). This pattern is broadly consistent with known functional differences across IPS, whereby anterior regions participate more strongly in sensorimotor transformations supporting reaching, grasping, and action planning, while posterior regions are more closely associated with visual-spatial processing (Duhamel et al., 1997, 1998; Snyder et al., 1997; Lewis and Van Essen, 2000a; Murata et al., 2000; Andersen and Buneo, 2002; Medendorp and Heed, 2019). Together, these findings suggest that IPS-S1 interactions are organized according to a large-scale topological hierarchy extending from the sulcal fundus toward the cortical convexities, rather than being distributed uniformly across IPS.

### Connectivity-defined transitions reveal functional heterogeneity within classical PPC areas

Interestingly, this organization showed partial but incomplete correspondence with classical cytoarchitectonic boundaries within PPC (Fig. 4, Fig. S4). Quantitative comparison with the Markov atlas confirmed that the functional and atlas-defined partitions shared systematic structure but were not related one-to-one. Instead, connectivity patterns were arranged primarily according to a seed group’s position within the sulcus and frequently extended across multiple cortical areas. This observation suggests that large-scale IPS-S1 connectivity may be constrained more strongly by continuous spatial organization across the sulcus than by discrete areal borders alone. Although functional organization within association cortex can be expressed as continuous gradients that coexist with, and sometimes transcend, classical cortical parcellations (Glasser et al., 2016; Margulies et al., 2016; Medendorp and Heed, 2019; Orban et al., 2021), the patchy connectivity patterns observed here indicate that IPS-S1 organization is not purely gradient-like. Within this framework, the semi-oval topology may represent a large-scale organizational scaffold related to sulcal geometry, within which more localized functional heterogeneity is expressed within and across individual PPC areas. The clustering analysis further revealed these local deviations from classical cytoarchitectonic organization. Several connectivity-defined clusters crossed established areal boundaries, whereas some individual cortical areas were subdivided into multiple connectivity compartments, producing patchy functional connectivity patterns (Fig. 4). Together, these findings suggest that IPS-S1 connectivity is organized across multiple spatial scales, combining a large-scale spatial scaffold structured by sulcal geometry with finer-scale, locally heterogeneous connectivity patterns within and across classical cortical areas.

The intermediate clusters were particularly informative in this regard. Positioned between the fundus cluster and the anterior convexity clusters, these transitional connectivity groups frequently aligned with the border between MIP and area 5 while simultaneously subdividing LIP into posterior-medial and anterior-lateral components (Fig. 4). However, individual-level boundary analyses showed that the reproducibility of these transitions was spatially heterogeneous (Fig. S5). Functional subdivisions within LIP were more consistently preserved across animals than those within VIP, area 5, and area 2, whereas the transition near the MIP– area 5 border showed comparatively greater inter-individual variability. Thus, the finer intermediate clusters should not necessarily be interpreted as fixed anatomical subdivisions shared identically across animals. Such observations indicate that IPS-S1 connectivity changes gradually across the cortical surface rather than abruptly at areal borders alone. This interpretation is also consistent with the systematic positional effects observed in the averaging analysis (Fig. 3), where connectivity varied progressively along both the anteroposterior and mediolateral dimensions of IPS. These findings suggest that IPS-S1 interactions cannot be described solely as a collection of discrete connectivity states associated with individual cortical areas. Instead, connectivity frequently changed in a graded manner across neighboring seed locations while still preserving localized specializations. In this respect, the present findings indicate that PPC organization contains both continuous and discrete features: large-scale connectivity transitions span the sulcus, whereas localized functional specializations remain associated with individual cortical areas. More broadly, this combination of continuous and localized patterns may indicate that functional organization in PPC emerges from multiple organizational principles that need not coincide with cytoarchitectonic boundaries. A similar principle has been proposed for motor cortex, where large-scale functional maps may arise from the interaction of multiple partially independent organizational dimensions rather than from a single anatomical map (Aflalo and Graziano, 2006), Such an organization could help explain why functional connectivity patterns do not always map directly onto anatomically defined cortical areas: localized functional specializations may reflect the combined influence of several connectivity or computational principles operating across the same cortical territory. From this perspective, the areal boundaries identified by cytoarchitectonic methods provide an important anatomical framework, but do not necessarily delimit functionally homogeneous units. The present findings therefore raise the possibility that the connectivity principles shaping IPS-S1 organization may constitute one component of a broader, multidimensional organization of PPC, potentially allowing somatosensory connectivity patterns to coexist with and interact with other functional systems, such as visual and auditory-related connectivity.

The repeated subdivision of LIP across S1 subregions and hemispheres is especially notable. Rather than exhibiting a single homogeneous connectivity profile, posterior-medial and anterior-lateral portions of LIP were consistently assigned to different connectivity clusters (Fig. 4). This observation is consistent with previous evidence that LIP contains substantial internal functional heterogeneity (Andersen et al., 1990), including distinct dorsal and ventral subdivisions identified by architectonic and connectivity-based studies (Lewis and Van Essen, 2000b). More broadly, this internal differentiation may be relevant to the greater subdivision of the human IPS, where multiple regions have been proposed to correspond to the macaque LIP complex (Kastner et al., 2017). Although this interpretation remains speculative, the present findings raise the possibility that the internal functional differentiation observed within macaque LIP reflects an organizational feature that can become further elaborated into multiple parietal regions across primate evolution. Previous studies have further suggested that posterior portions of LIP are more strongly associated with visual-spatial and retinotopic processing, whereas anterior regions exhibit greater sensorimotor and multisensory integration (Blatt et al., 1990; Ben Hamed et al., 2001; Sereno et al., 2001; Ben Hamed and Duhamel, 2002; Premereur et al., 2011). The present results extend this view by suggesting that these functional differences may also be reflected in distinct patterns of interaction with somatosensory cortex.

Together, these results indicate that IPS-S1 connectivity reflects both continuous spatial organization across the sulcus and localized functional specialization within individual PPC areas.

### Heterogeneous somatosensory input contributes to functional diversity among IPS-S1 connectivity clusters

The four S1 subregions differ not only in the modality of their dominant thalamic input but also, within each subregion, in the specific body part represented at any given cortical location (Kaas et al., 1979; present results, Fig. 5). Consistent with this organization, the somatotopic composition analysis in Fig. 6 showed that connectivity-defined IPS clusters were generally associated with combinations of putative body-part sectors rather than with a single somatotopic representation, and that these combinations varied across S1 subregions and IPS clusters. Any IPS territory functionally coupled with S1 therefore can be characterized along two largely independent axes of heterogeneity: the modality/processing stream with which it is coupled (i.e. which S1 subregion), and the body-part identity associated with that connectivity (i.e. which medio-lateral part of the S1 subregion).

This dual heterogeneity offers a candidate explanation for why the connectivity-defined clusters identified here (Fig. 4) do not map cleanly onto single cytoarchitectonic areas or single body-part zones, but instead cut across both. A cluster more strongly coupled with area 2 versus area 3a input would be preferentially associated with multimodal somatosensory processing rather than predominantly proprioceptive processing, regardless of which body part is represented; conversely, two clusters both showing prominent connectivity with area 2 input but differing in their relative connectivity with different somatotopic zones, for instance hand versus face, would share a similar S1 processing stream while emphasizing functionally distinct effectors. The heterogeneous body-part profiles observed across the connectivity-defined clusters in Fig. 6 are consistent with this combination of S1-subregion and somatotopic variation. The repeated subdivision of LIP into posterior-medial and anterior-lateral connectivity compartments, and the intermediate edge-layer clusters spanning the MIP/area 5 border, are consistent with IPS territories combining these two sources of variation in different proportions rather than exhibiting connectivity with a single homogeneous component of S1.

### Somatotopic boundaries revealed by connectivity minima

Local minima in the connectivity profiles aligned with previously reported somatotopic boundaries (Fig. S6), suggesting that large-scale network architecture preserves aspects of body-part organization traditionally identified using electrophysiological recordings and sensory localizers (Kaas et al., 1979; Nelson et al., 1980; Pons et al., 1985; Arcaro et al., 2019). These observations suggest that neighboring somatotopic territories participate in partially distinct patterns of corticocortical interaction and that transitions between body-part representations can therefore emerge as inflection points within connectivity space.

Notably, some somatotopic boundaries were reflected more strongly than others. Connectivity-derived minima corresponded most consistently to the putative tongue-face and hand/trunk-leg transitions, whereas the classical face-hand boundary was less robustly identified across S1 subregions and hemispheres. One possible explanation is that different somatotopic boundaries are associated with different degrees of functional and anatomical separation. Because functional connectivity reflects coordinated activity between distributed cortical territories, functionally related body-part representations might be expected to exhibit less sharply differentiated connectivity patterns than representations that participate in more distinct functional networks. This possibility is consistent with proposals that hand and face representations may be particularly strongly integrated in primates because of the importance of coordinated hand-mouth actions (Graziano and Aflalo, 2007). Consistent with this possibility, intrinsic anatomical connectivity near the face-hand transition is not uniformly segregated across all lower-face representations. Manger et al. (1997) showed that upper lip and muzzle representations exhibit little overlap in intrinsic horizontal connections with the hand representation, whereas the lower jaw/neck representation remains strongly interconnected with hand territory. Such partial continuity in anatomical connectivity near the lower-face region may produce smoother large-scale IPS-S1 connectivity transitions and thereby reduce the prominence or spatial consistency of connectivity-derived minima near portions of the face-hand interface.

Interestingly, different putative boundaries were not recovered uniformly across S1 subregions. Tongue-face transitions were most evident in areas 3a and 3b, whereas hand/trunk-leg transitions were observed more consistently in areas 1 and 2. Although the present study was not designed to determine the origin of these differences, they likely reflect known functional distinctions among S1 subregions. Areas 3a, 3b, 1, and 2 differ in their receptive-field properties and relative contributions to proprioceptive and tactile processing (Kaas et al., 1979; Delhaye et al., 2018), leading to differences in how somatotopic information is represented within IPS-S1 interaction networks.

Importantly, connectivity-derived minima should not be interpreted as direct substitutes for electrophysiological mapping or functional localizers. Functional connectivity measures large-scale correlations in spontaneous cortical activity rather than stimulus-evoked neuronal responses. Nevertheless, the correspondence between connectivity minima and independently established somatotopic boundaries suggests that intrinsic cortical dynamics can preserve features of somatosensory organization at the level of large-scale connectivity. The extent to which individual minima are reproducible across datasets, sampling schemes, or analytical choices remains an important consideration when interpreting these features. Functional connectivity may therefore provide a complementary systems-level perspective on S1 architecture by revealing how body-part representations are embedded within distributed cortical interaction networks, while the present minima should be regarded as candidate connectivity signatures of somatotopic transitions rather than definitive markers of cortical boundaries.

### Methodological significance, limitations, and future directions

The present study highlights the utility of dense spatial sampling combined with resting-state functional connectivity for investigating cortical organization. An important strength of the dataset is that recordings were obtained from awake macaques and included a relatively large number of animals (n = 10). Because anesthesia can substantially alter functional connectivity patterns and network architecture (Vincent et al., 2007; Grandjean et al., 2014; Hutchison and Everling, 2014; Xu et al., 2018; Thomas et al., 2021; Manickam et al., 2026), awake-state measurements provide a closer approximation to physiological network organization. Furthermore, the sample size enabled group-level analyses despite the fine spatial resolution of the seed-based approach. Rather than restricting analyses to a small number of predefined regions of interest, we systematically sampled connectivity across an IPS-centered PPC territory and examined how connectivity patterns changed as a continuous function of spatial position. This approach enabled the identification of organizational features that might be difficult to detect using conventional area-based analyses, including the semi-oval topology, gradual connectivity gradients across the sulcus, and connectivity-defined transitions associated with somatotopic boundaries.

To emphasize the spatial organization of IPS-S1 interactions, subsequent analyses focused on the prevalence of positive connectivity rather than absolute connectivity strength. By quantifying the spatial distribution of connected locations across IPS and S1, this representation highlights large-scale topological relationships while reducing the influence of localized variations in connectivity magnitude. Previous work using the same awake macaque dataset demonstrated that the proportion of significantly connected surface voxels provides a sensitive measure of large-scale functional organization when comparing connectivity patterns across extensive cortical territories (Sheng et al., 2025). By treating connectivity as a spatially varying phenomenon rather than as a set of pairwise relationships between predefined areas, the present framework provides a complementary perspective on cortical organization that bridges traditional areal parcellations and large-scale network analyses (Hutchison and Everling, 2014; Haak et al., 2018). Similar dense-sampling approaches may prove useful for investigating organizational principles in other association cortices where functional boundaries are not easily captured by existing anatomical definitions.

Several limitations should nevertheless be considered. First, resting-state functional connectivity measures statistical relationships in spontaneous activity and therefore cannot establish the directionality or causal basis of interactions between IPS and S1. Consequently, the connectivity-defined subdivisions identified here should not be interpreted as direct anatomical pathways or functional circuits. Second, although the observed connectivity patterns showed substantial correspondence with somatotopic organization, and only partial correspondence with classical cytoarchitectonic boundaries, the present analyses were performed using resting-state data acquired in the absence of explicit sensory or motor behavior. Future studies combining dense connectivity sampling with task-based paradigms, electrophysiological recordings, or anatomical tracing could help clarify the functional significance of the identified connectivity gradients and cluster boundaries. Such multimodal approaches have already proven valuable for relating resting-state functional connectivity to anatomical and physiological organization in the macaque brain (Shen et al., 2012; Howells et al., 2020; Tang et al., 2025). In particular, determining how the semi-oval topology relates to sensory receptive fields, body-part representations, and sensorimotor transformations within IPS will be important for understanding the mechanisms that give rise to the large-scale connectivity organization observed here.

## Data and code availability

Data and code are available upon request.

## Author contributions

Conceptualization: W.-A. S. and S.B.H.; Data Curation: W.V., W.-A. S., and M.F.; Formal Analysis: W.-A. S. and S.B.H.; Funding Acquisition: S.B.H.; Investigation: W.-A. S., and S.B.H.; Methodology: W.-A. S., S.C., M.F., and S.B.H.; Resources: W.V.; Supervision: S.B.H.; Validation: S.B.H.; Visualization: W.-A. S., C.S.; Writing—Original draft: W.-A. S. and S.B.H.; and Writing—review & editing: W.-A. S., S.C., M.F., W.V., T.H., and S.B.H.

## Ethical statement

Animal care and experimental procedures were performed in accordance with the National Institute of Health’s Guide for the Care and Use of Laboratory Animals, the European legislation (Directive 2010/63/EU) and were approved by the Animal Ethics Committee of the KU Leuven. Weatherall reports were used as reference for animal housing and handling. All animals were group-housed in cages sized 16–32 m3, which encourages social interactions and locomotor behavior. The environment was enriched by foraging devices and toys. The animals were fed daily with standard primate chow supplemented with fruits, vegetables, bread, peanuts, cashew nuts, raisins, and dry apricots. The animals were exposed to natural light and additional artificial light for 12 h every day. On training and experimental days, the animals were allowed unlimited access to fluid through their performance during the experiments. Using operant conditioning techniques with positive reinforcers, the animals received fluid rewards for every correctly performed trial. During non-working days, they received water in their living quarters. Throughout the study, the animals’ psychological and veterinary welfare was monitored daily by the veterinarians, the animal facility staff, and the lab’s scientists, all specialized in working with non-human primates. The animals were healthy at the conclusion of our study.

## Acknowledgments

We also thank Thomas Perret, Johan Pacquit, and Marco Bimbi for help with the hardware computational resources.

## Supplementary figures

**Supplementary Figure 1.**
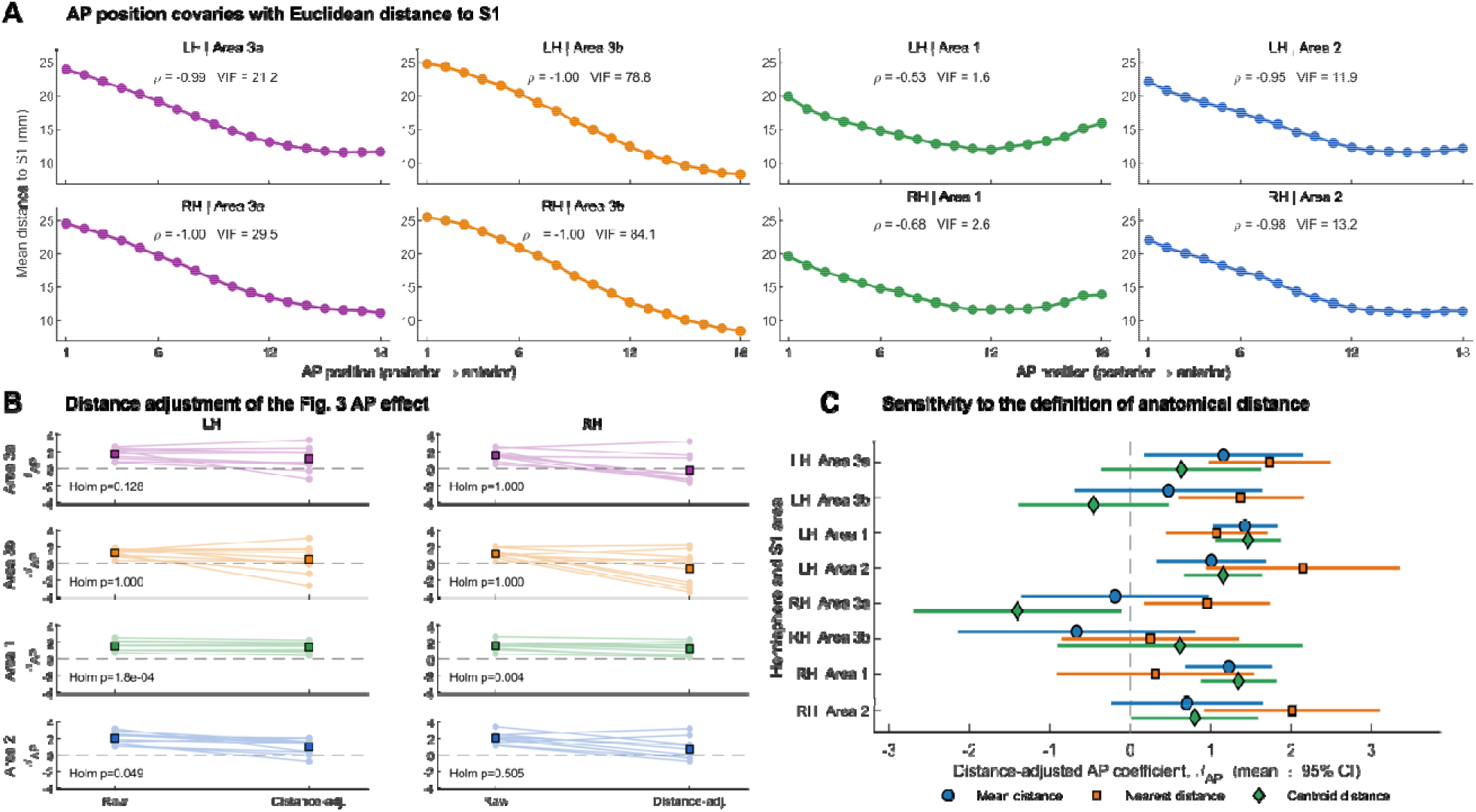
Influence of anatomical distance on the anteroposterior IPS-S1 connectivity gradient. **A.** Relationship between IPS anteroposterior position and Euclidean distance to each S1 subregion. AP1 corresponds to the posterior IPS and AP18 to the anterior IPS. For each AP position, mean Euclidean distance was calculated across the same IPS seed positions used for the Fig. 3 anteroposterior analysis (m5–l5; k = 3:13). Distances were calculated from each IPS seed to all surface vertices within the corresponding S1 subregion and then averaged across seeds at each AP position. Spearman’s ρ and the variance inflation factor (VIF) quantify the association and collinearity between AP position and distance. **B.** Effect of controlling for Euclidean distance on the Fig. 3 anteroposterior connectivity gradient. For each monkey (n = 10), an AP coefficient β_AP_ was estimated from the Fig. 3-matched connectivity profiles before and after including mean Euclidean distance to S1 as a covariate. Positive β_AP_ indicates stronger connectivity toward the anterior IPS. Lines connect estimates from the same monkey; colored squares indicate group means. Holm-adjusted p values are shown for the distance-adjusted AP coefficients across the eight hemisphere × S1-subregion comparisons. The AP effect remained robust in area 1 bilaterally and in left area 2, whereas stronger AP–distance collinearity was present particularly in areas 3a and 3b. **C**. Sensitivity of the distance-adjusted AP effect to the definition of anatomical distance. Points show the mean β_AP_ across monkeys and horizontal lines indicate 95% confidence intervals for models using mean distance to all S1 vertices, distance to the nearest S1 vertex, or distance to the S1 centroid. The vertical dashed line indicates β_AP_ = 0.

**Supplementary Figure 2.**
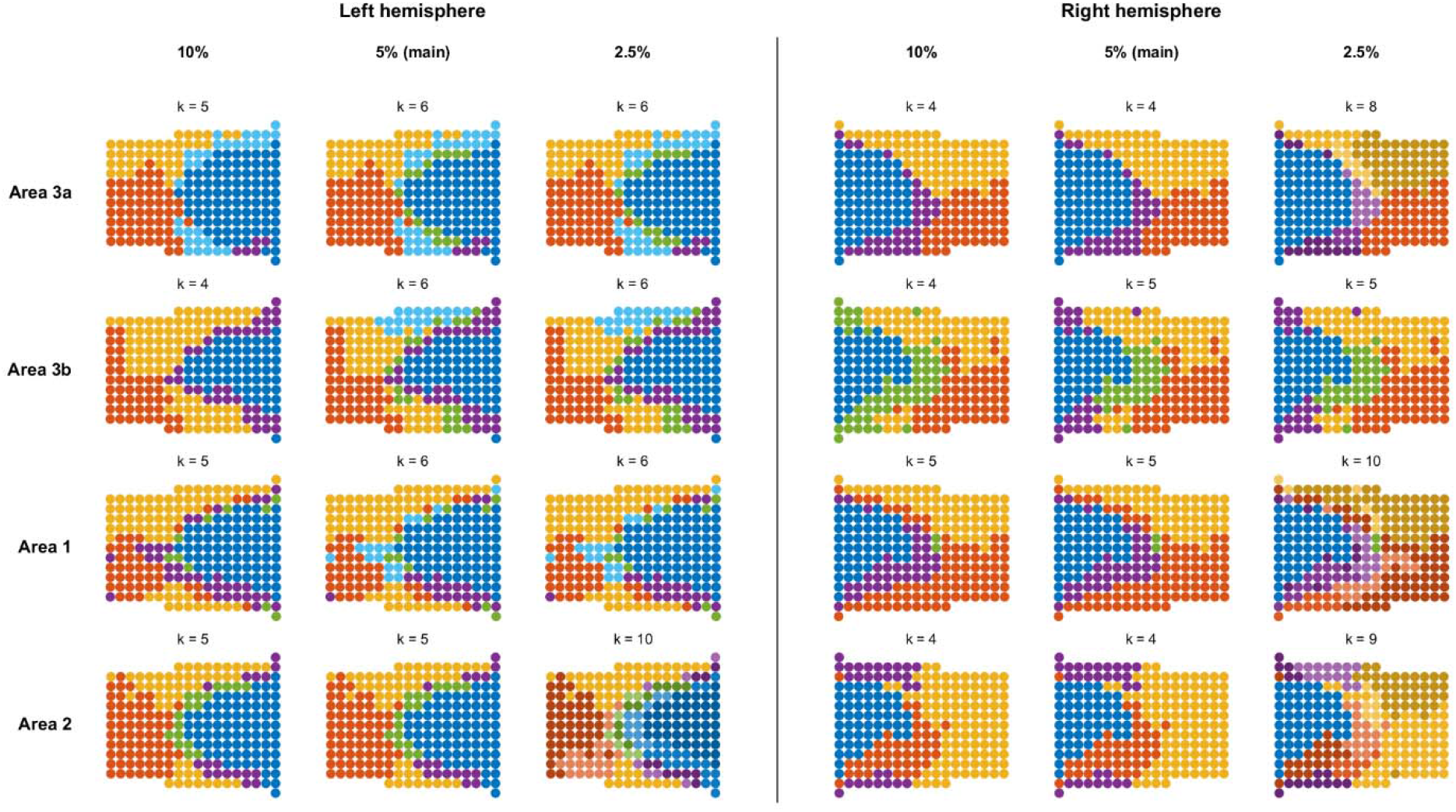
Robustness of IPS connectivity-defined clustering to the cluster-size criterion. Spatial distributions of IPS seed clusters are shown separately for the left and right hemispheres and for S1 areas 3a, 3b, 1, and 2. Hierarchical clustering was repeated using the same connectivity data, Chebyshev distance metric, average-linkage procedure, and iterative cluster-selection rule as in the primary analysis, while varying only the cluster-size threshold from 10% to 5% to 2.5% of IPS seeds. The 5% condition corresponds to the criterion used in the main analysis, whereas 10% and 2.5% provide more conservative and more permissive clustering resolutions, respectively. Each circle represents one IPS seed, and the selected number of clusters (k) is indicated above each map. To facilitate comparison across clustering resolutions, colors were referenced to the 5% solution: clusters at 10% and 2.5% were matched to the 5% cluster with which they shared the greatest seed overlap, and finer subdivisions of the same 5% cluster were displayed using related shades of the same base color. This analysis assesses whether the large-scale spatial organization of IPS connectivity-defined territories is preserved across reasonable choices of the cluster-size threshold.

**Supplementary Figure 3.**
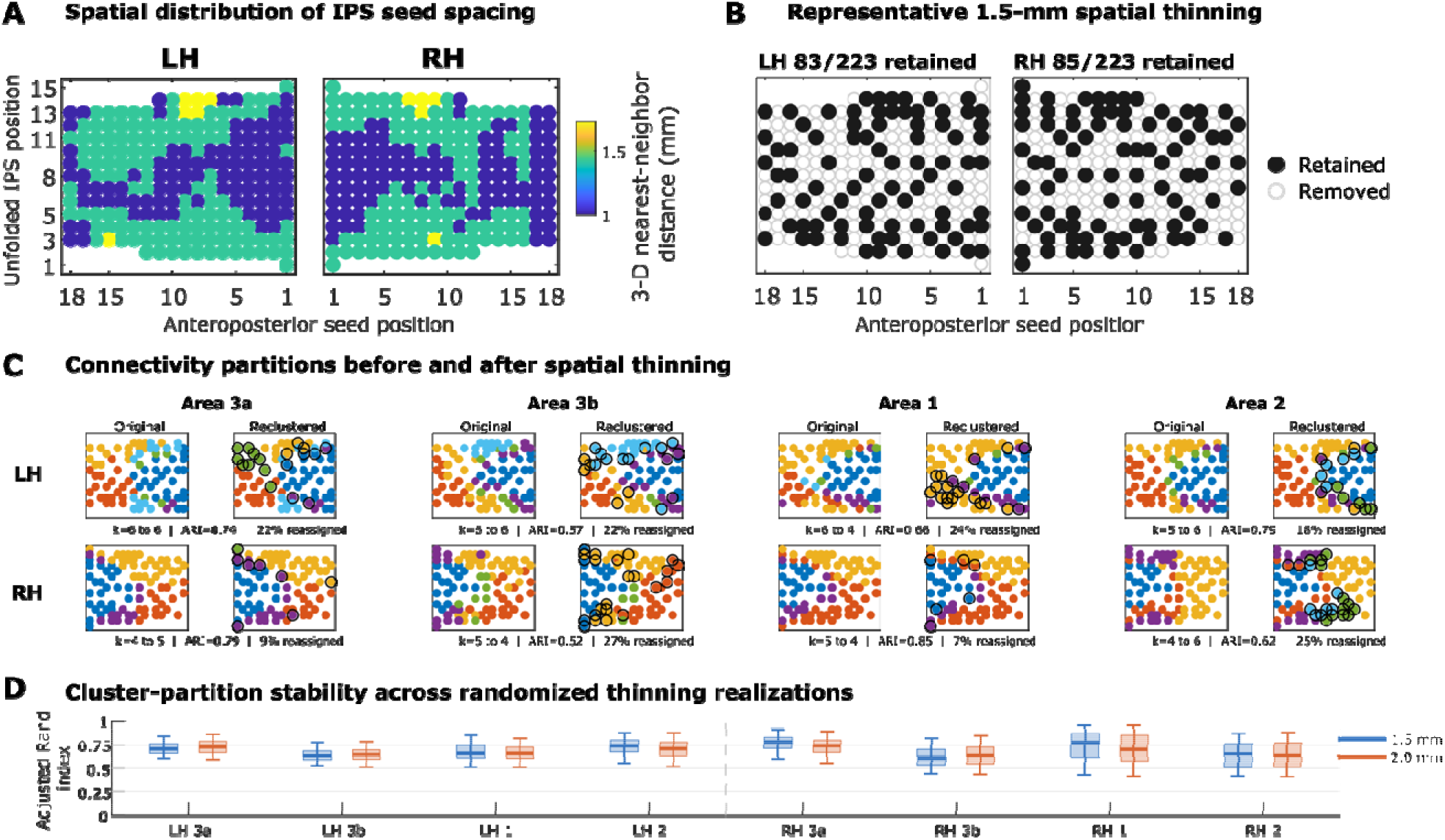
Robustness of connectivity-defined IPS organization to non-uniform spatial sampling. A. Spatial distribution of the three-dimensional nearest-neighbor distance among the 223 IPS seed voxels in the left (LH) and right (RH) hemispheres. Shorter nearest-neighbor distances indicate regions of denser sampling. B. Representative spatial-thinning realizations using a minimum inter-seed separation of 1.5 mm. Retained seeds are shown as filled circles and removed seeds as open circles. The representative realizations retained 83 of 223 seeds in LH and 85 of 223 seeds in RH. C. Comparison of the original connectivity-defined cluster assignments with those obtained after reclustering the representative spatially thinned seed sets, shown separately for S1 areas 3a, 3b, 1, and 2. For each pair, the same retained seeds are displayed in the original and reclustered maps. Reclustering colors were matched to the original clusters by maximum overlap for visualization only. Black outlines indicate retained seeds whose cluster assignment changed after reclustering. The number of clusters, adjusted Rand index (ARI), and proportion of reassigned seeds are indicated below each pair. D. Distribution of ARI values across 500 randomized spatial-thinning realizations for minimum inter-seed separations of 1.5 and 2.0 mm. Boxes indicate the interquartile range, horizontal lines indicate the median, and whiskers indicate the 5th–95th percentile range. Despite substantial reduction in the number of retained seeds, the major connectivity-defined partition structure was largely preserved, whereas the exact number and boundaries of finer subdivisions showed greater variability.

**Supplementary Figure 4.**
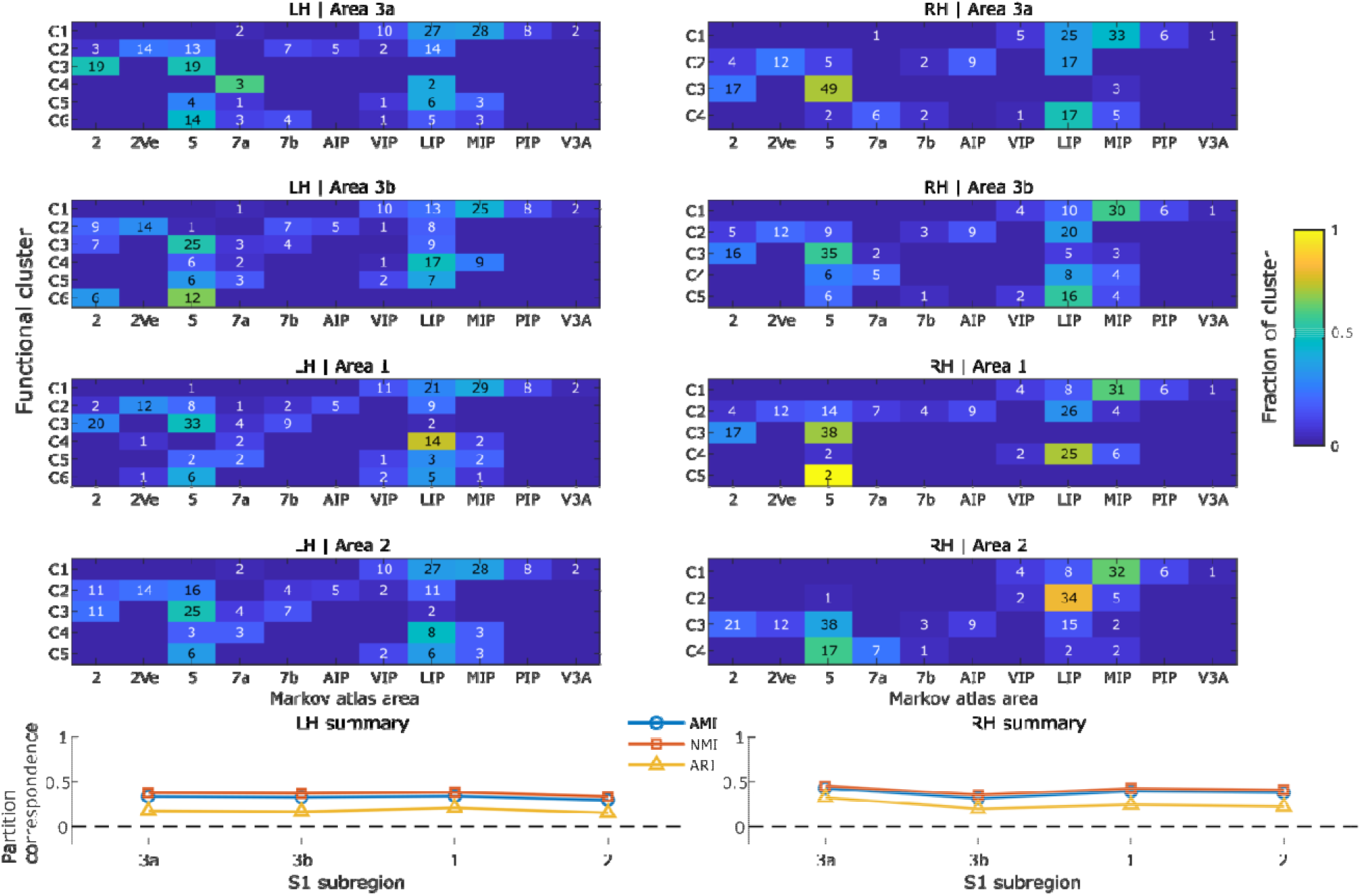
Correspondence between connectivity-defined IPS clusters and Markov atlas areas. For each S1 subregion (areas 3a, 3b, 1, and 2), connectivity-defined IPS clusters were compared with the digitized Markov atlas assignments of the 223 IPS seed positions. Heatmaps show the anatomical composition of each functional cluster separately for the left (LH) and right (RH) hemispheres. Columns indicate Markov atlas areas and rows indicate functional clusters. Color represents the fraction of seeds within each functional cluster assigned to the corresponding atlas area, and numbers indicate the absolute number of seeds in each cell. One unlabeled seed in the right hemisphere was excluded from the correspondence analysis. Bottom panels summarize partition similarity using adjusted mutual information (AMI), normalized mutual information (NMI), and adjusted Rand index (ARI). Higher values indicate greater correspondence between the connectivity-defined and cytoarchitectonic partitions. Across S1 subregions, the analyses revealed partial but incomplete correspondence between functional clustering and Markov atlas organization.

**Supplementary Figure 5.**
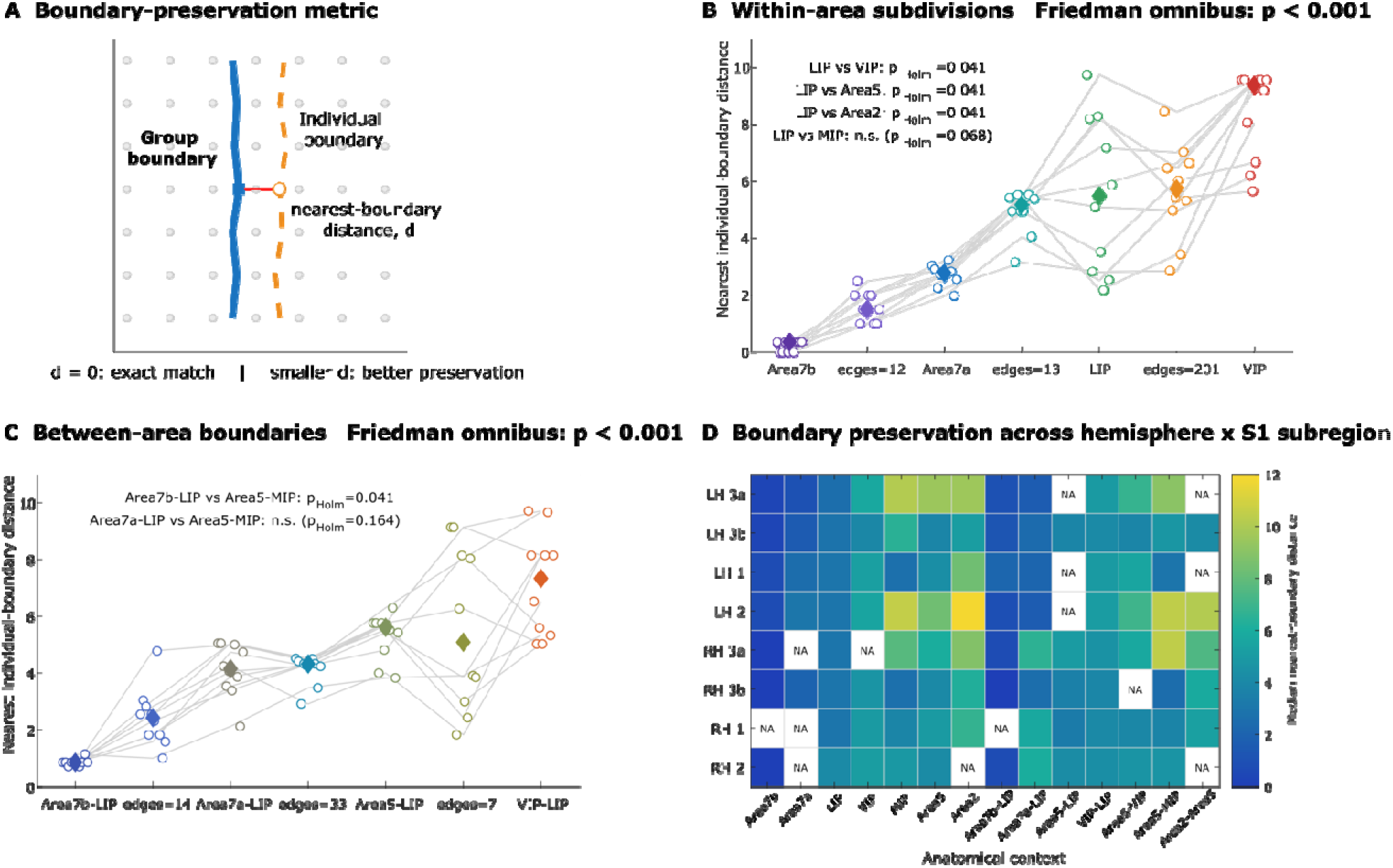
Individual-level reproducibility of group-defined IPS functional boundaries varies across cytoarchitectonic territories. **A.** Schematic of the boundary-preservation metric. For each group-defined functional boundary edge, the distance to the nearest boundary edge in the corresponding individual-animal fixed-k clustering solution was measured on the unfolded IPS lattice. A distance of 0 indicates an exact edge match, whereas smaller non-zero values indicate a nearby spatially displaced boundary. Lower nearest-boundary distance therefore indicates better preservation of the group-defined boundary in individual animals. **B.** Reproducibility of functional subdivisions located within the same Markov atlas area. For each monkey, nearest-boundary distances were first summarized within each hemisphere × S1-subregion block and then across available blocks, yielding one monkey-level value per anatomical context. Circles indicate individual monkeys and diamonds indicate the median across monkeys. The number of distinct group-level boundary edges contributing to each anatomical context is shown below each label. Boundary preservation differed significantly across within-area contexts (Friedman test, Holm-corrected across the two predefined analysis families, p < 0.001). Selected post-hoc comparisons involving LIP are shown in the panel; p values were Holm-corrected across all within-area pairwise comparisons. **C.** Reproducibility of functional boundaries located between pairs of Markov atlas areas. Values were summarized using the same monkey-balanced procedure as in panel B. Boundary preservation differed significantly across between-area contexts (Friedman test, Holm-corrected across the two predefined analysis families, p < 0.001). Selected post-hoc comparisons are shown in the panel; p values were Holm-corrected across all between-area pairwise comparisons. Boundaries involving the lateral convexity, particularly 7b–LIP and 7a–LIP, showed relatively low nearest-boundary distances, whereas boundaries such as area 5–MIP and area 2–area 5 showed poorer individual-level preservation. **D.** Boundary preservation shown separately for each hemisphere and S1 subregion. Each heatmap cell was calculated by first taking the median nearest-boundary distance across relevant group-boundary edges within each monkey and then taking the median across monkeys. Lower values indicate better preservation. NA cells indicate anatomical boundary contexts that were not represented in the corresponding group-level hemisphere × S1-subregion clustering and therefore could not be evaluated. Only anatomical contexts represented by at least five distinct group-level boundary edges and available in all 10 monkeys were included in the inferential analyses.

**Supplementary Figure 6.**
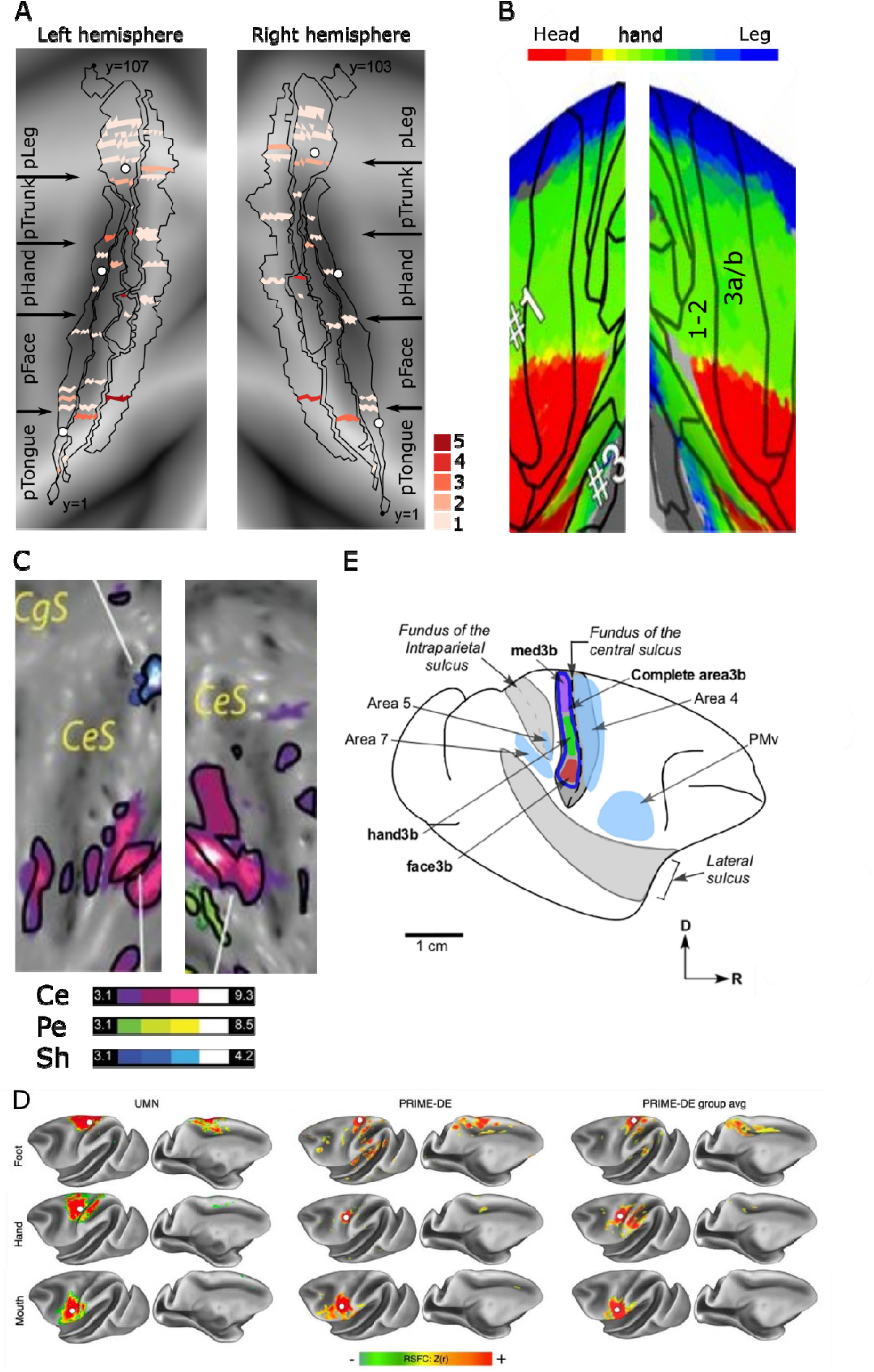
Positive-connectivity local minima and somatotopic functional localizers. **A.** Local minima identified from connectivity clusters in areas 3a, 3b, 1, and 2. Same as figure 5. **B.** Cropped and adapted from Arcaro et al. (2019), showing group-average body maps from two juvenile macaques. Contralateral representations of the face, hand, and foot are distributed throughout the anterior parietal cortex. Data were overlaid onto the Saleem and Logothetis atlas and thresholded at a group-average z-statistic > 4 for the strongest independent component (IC) assigned to each vertex. **C.** Cropped and adapted from Wardak et al. (2016), showing cortical regions preferentially responsive to the face center (Ce, purple; t ≥ 3.1, p < 0.001, uncorrected) relative to the face periphery (Pe, green) and shoulder (Sh, blue) representations. Black outlines indicate regions selectively representing one of the body part stimulations. CeS, central sulcus; CgS, cingulate sulcus. **D.** Functional connectivity maps seeded from a continuous line of points down anterior central sulcus (rows 1-3), in fMRI data from an individual macaque scanned for 77 min on a 10.5T MRI scanner (left); an individual macaque scanned for 53 min on a 3T scanner (middle); and group-averaged data from eight macaques each scanned for 53 min on a 3T scanner (right). These seeds demonstrated connectivity patterns corresponding to the known functional divisions between M1 regions representing the foot (second row), hand (third row), and face (bottom row). From Gordon et al., 2023. **E.** Schematic representation of area 3b ROI (thick dark blue outline; area3b), and ROIs of different body part representations: face, face3b (red); hand, hand3b (green); and medial body, med3b (violet), from Thomas et al., 2021.

**Supplementary Figure 7.**
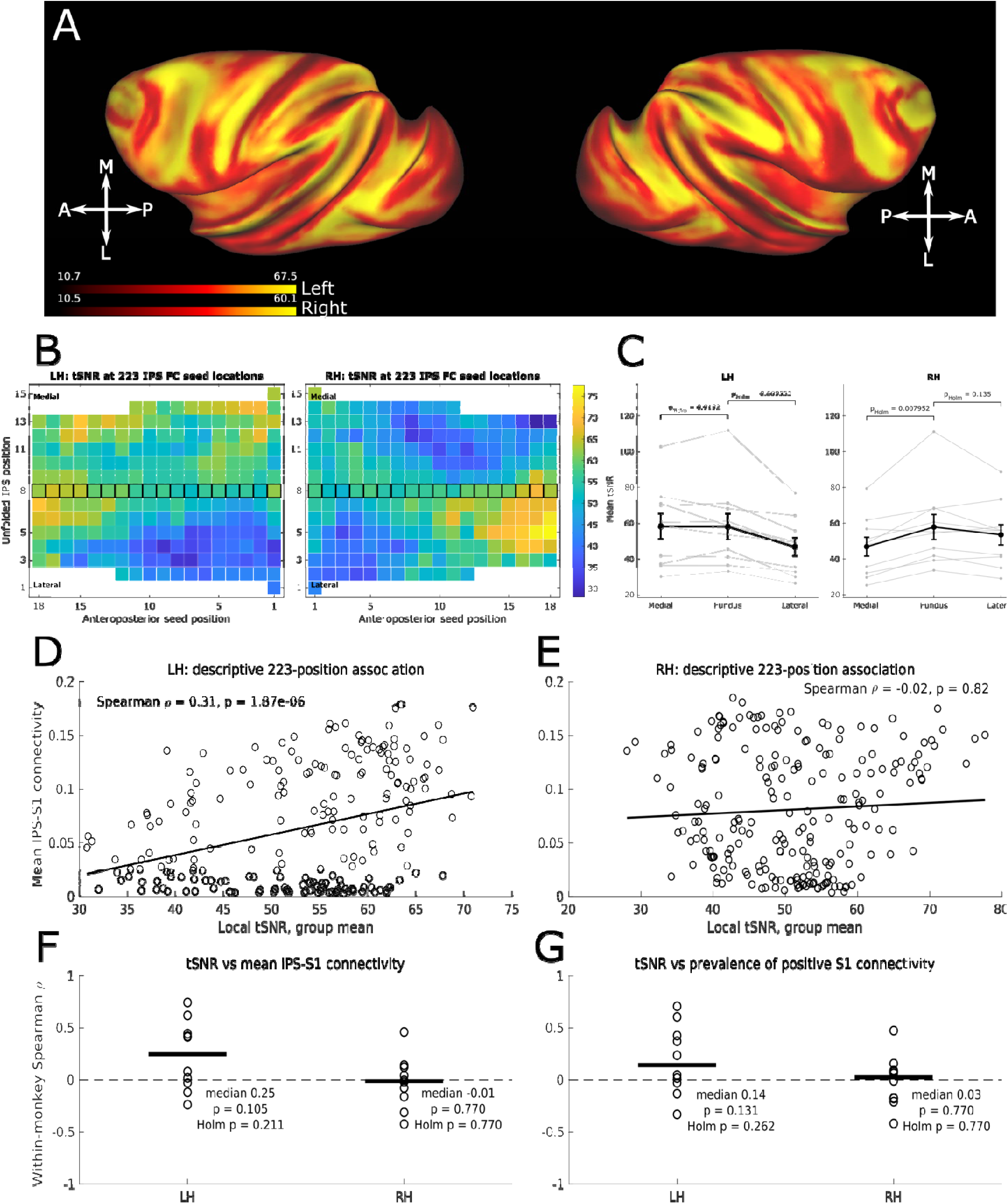
Temporal signal-to-noise ratio across the intraparietal sulcus. **A.** Group-average temporal signal-to-noise ratio (tSNR) projected onto the cortical surface for the left and right hemispheres. Hemisphere-specific color scales were used for surface visualization. **B.** Group-average tSNR sampled at the 223 single-voxel IPS seed locations used for the functional connectivity analyses in each hemisphere. Seed values are displayed using the same unfolded IPS organization used to represent IPS connectivity, with medial and lateral positions shown along the vertical axis and anteroposterior seed position along the horizontal axis. The fundus seed row is outlined in black. A common color scale is used for both hemispheres. **C.** Mean tSNR across medial, fundal, and lateral IPS seed groups for each monkey. Thin gray lines represent individual monkeys, and black symbols and error bars indicate the group mean ± SEM. Fundal tSNR was comparable to medial IPS and higher than lateral IPS in the left hemisphere (Holm-corrected paired comparisons: fundus vs. medial, p = 0.9132; fundus vs. lateral, p = 0.0096), whereas in the right hemisphere fundal tSNR was higher than medial IPS and comparable to lateral IPS (fundus vs. medial, p = 0.0080; fundus vs. lateral, p = 0.135). Thus, the reduced IPS–S1 functional connectivity observed near the fundus was not accompanied by a corresponding reduction in local tSNR. **D-G.** Relationship between local temporal signal-to-noise ratio (tSNR) and mean IPS–S1 functional connectivity across the 223 IPS seed positions in the left (D) and right (E) hemispheres. Each point represents one IPS seed position, with tSNR and functional connectivity averaged across monkeys. Mean functional connectivity was calculated across structurally valid mediolateral bins from S1 areas 3a, 3b, 1, and 2. These group-average correlations are shown for visualization only because neighboring IPS seed positions are not statistically independent. **F.** Within each monkey, Spearman correlations were calculated across the 223 IPS seed positions between local tSNR and mean IPS–S1 functional connectivity. Points represent individual monkeys (n = 10), and horizontal bars indicate the median correlation coefficient. Correlation coefficients were tested against zero using two-sided Wilcoxon signed-rank tests; Holm-corrected p values account for the two hemispheres. **G.** Same analysis using the prevalence of positive S1 connectivity, defined as the proportion of structurally valid S1 bins with positive functional connectivity (>0), providing a measure consistent with the binarized connectivity representation used in the main analyses. Neither measure showed a consistent positive association with local tSNR across monkeys.

